# Integration of Audiovisual Danger Stimuli in the Zebrafish Tectum is Linked with Premotor and Behavioral Enhancements

**DOI:** 10.64898/2025.12.30.697023

**Authors:** Nicolás Martorell, Violeta Medan

## Abstract

Multisensory integration enables animals to combine sensory cues, enhancing threat detection and escape behaviors. Although the Tectum is traditionally considered a visual center, its role in integrating auditory and visual danger signals is not fully understood. Using in vivo calcium imaging in larval zebrafish, we demonstrate that auditory stimuli recruit a significant population of tectal neurons, mainly in deep layers. Visual looms elicit strong tectal responses, but combined auditory-visual stimuli robustly increase neural activity and the likelihood of rapid escape, especially for less salient cues engaging more tectal neurons than unisensory stimuli. These results establish the zebrafish Tectum as a multisensory processing area. Notably, the heightened tectal response during multisensory stimulation is strongly correlated with increased activity in hindbrain neurons implicated in escape-like motion events, connecting tectal integration to premotor circuits that drive behavioral output. Our findings provide a mechanistic basis for tectal multisensory processing in enhancing behavioral escape responses.

## INTRODUCTION

Multisensory integration is a neural process by which animals combine information from different sensory modalities to build a coherent representation of their environment^1,2^. This capability is essential for reducing ambiguity of sensory inputs, such as visual, auditory, and tactile cues, enhancing an organism’s ability to detect and react to critical stimuli, such as predators, prey, or environmental hazards^3–6^.

This adaptive process has been widely documented across various animal species and contexts^7–10^. In the mammalian superior colliculus, neurons integrate multisensory stimuli due to synaptic convergence of afferents conveying information from different sensory modalities^11^. Early studies in cats established that the neural response to multisensory stimulation can be supralinear with respect to the sum of the unisensory responses, and that integration effects are larger when weak or ambiguous stimuli are combined^12,13^.

In other vertebrates, the Tectum, a structure homologous to the superior colliculus, is known to receive inputs from multiple sensory modalities^14–18^. This brain region is organized into distinct layers, primarily the neuropil-which receives afferents from retinal and other structures- and the periventricular layer (PVL) composed of tectal somata^15,17,19^. Retinotopic inputs of the visual field are received in the superficial layers of the neuropil, while deeper layers receive inputs from additional sensory modalities, like auditory, somatosensory and lateral-line inputs. Neurons in the PVL are known to project to downstream targets in the hindbrain and spinal cord, where premotor neurons are involved in selecting and executing motor responses^20–24^. Despite receiving inputs from multiple sensory modalities and projecting to downstream motor centers, it is still unknown if the Tectum performs multisensory integration and if that integration influences motor output.

The Tectum of zebrafish (*Danio rerio*) has been traditionally associated with visual processing^25,26^. While a few studies showed that tectal neurons can respond to non-visual stimuli such as auditory and water flow cues, responses were weak and sparse^27–29^ and one study reported multisensory suppression in some tectal neurons^26^. However, no reports of robust multisensory enhancement in the zebrafish Tectum have been reported. Furthermore, it is not known whether tectal integration, if present, plays a functional role in influencing behavioral outputs.

Zebrafish are vision and hearing specialists with a well documented behavioral repertoire^30,31^, making it a valuable model for studying audiovisual integration^25,28,32^. Specifically, danger stimuli evoke rapid escape responses known as C-starts, which allow fish to evade potential threats with remarkable speed and precision^33^. These fast escape responses are known to be triggered by visual stimuli (e.g., looming objects) and auditory cues (e.g. abrupt sounds)^34–36^. Visual information reaches the tectum from the retinal ganglion cells where it is conveyed to downstream circuits. In larval zebrafish auditory information is principally encoded by movement of saccular and utricular otoliths and is conveyed by the VIIIth nerve afferents through the octavolateralis nuclei in the hindbrain to the torus semicircularis (inferior colliculus in mammals) in the midbrain, and then to the thalamus^37^. Two types of fast escapes have received considerable attention due to their distinctive motor pattern and identifiable neural basis. The fastest escapes are short-latency C-starts (SLC), which are mediated by a pair of hindbrain premotor neurons, the Mauthner cells, which receive direct auditory input from 8th nerve afferents and indirect visual input through tectal projections^22,38,39^. Long-latency C-starts (LLC) are slower than SLC and mediated by Mauthner cell homologs and other hindbrain circuits^40–44^ but their sensory inputs are less clear.

In this study, we investigate whether audiovisual integration occurs within the Tectum and characterize its effects on behavioral decision-making in larval zebrafish. Through behavioral essays and calcium imaging at single-cell resolution we describe a strong multisensory enhancement of neural activation in the Tectum and of C-start probability during escape behavior. We observe robust auditory responses in the deep layers of the Tectum, extending its role in sensory processing beyond visual computation. Moreover, we find that integration in the tectum is particularly effective when visual stimuli are harder to detect and that a fraction of tectal neurons show supralinear responses to multisensory stimulation. Crucially, we find that tectal multisensory enhancement is strongly correlated with the activation of premotor neurons in the hindbrain, providing evidence for the zebrafish Tectum is a key site for multisensory integration and behavior selection in the context of threat detection.

## RESULTS

### Multisensory Stimulation Enhances Escape Probability in Freely-Swimming Fish

To determine if multisensory information influences escape behaviors of larval zebrafish, we conducted behavioral experiments in 4-7 dpf freely swimming zebrafish (n = 40, Fig. 1A). Visual and auditory stimuli of different intensities were presented individually and in combination, including three levels of visual contrast, three levels of auditory amplitude, and their nine combinations, along with a no-stimulus control (Fig. 1B-C). For these experiments we designed a multisensory stimulus consisting of a looming stimulus (with fixed dynamics and variable contrast) which expands until on its maximum diameter a short and abrupt acoustic stimulus (of variable intensity) is presented. This stimulus configuration aims to mimic the approaching of a predator that can be seen from some distance but that produces a strong vibrational stimulus when it is very close and about to strike.

**Figure 1.**
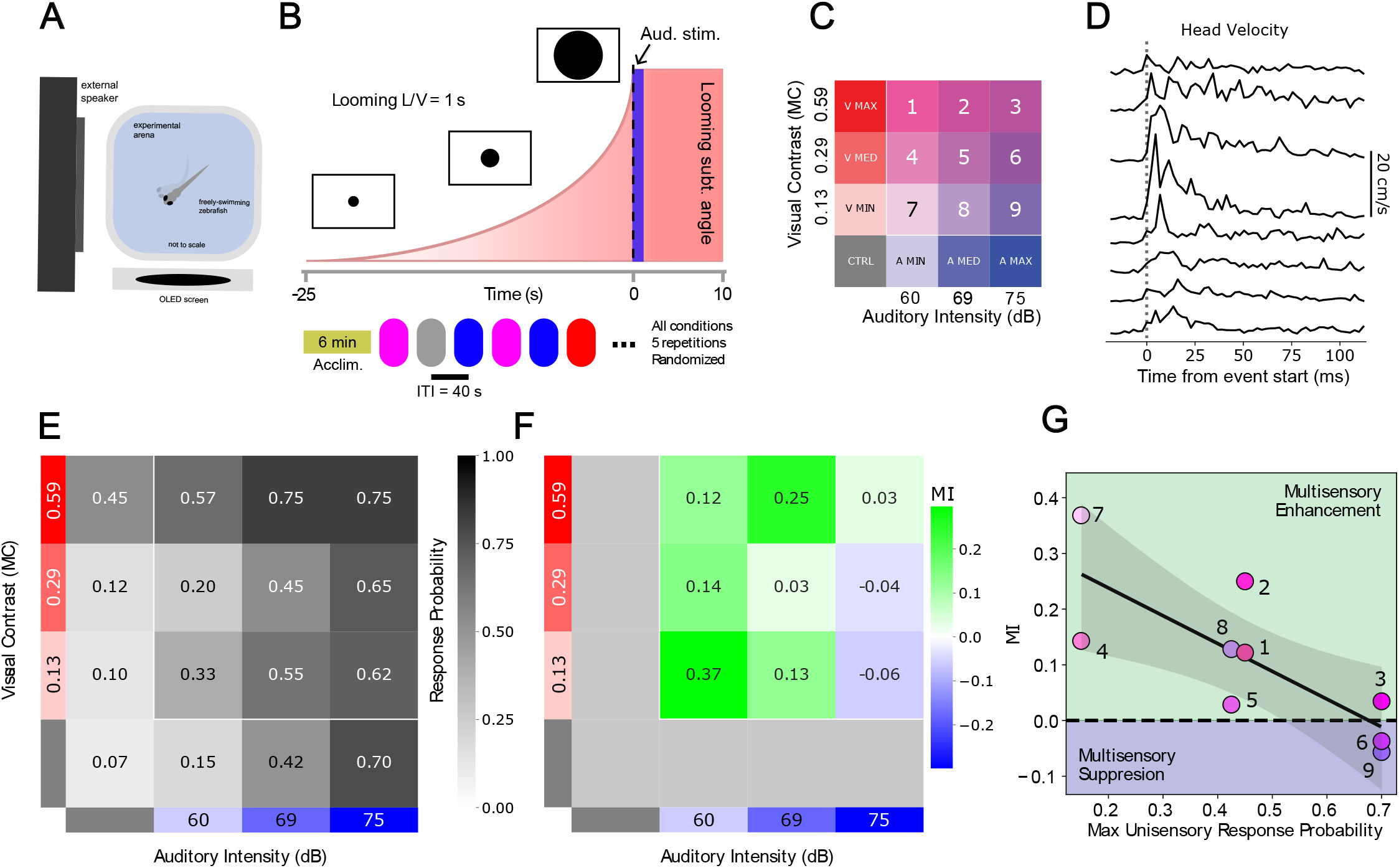
Multisensory enhancement of escape probability is inversely related to stimulus intensity. (A) Experimental arena used to present visual and auditory stimuli to freely-swimming zebrafish (N = 40). (B) Looming stimulus dynamics (l/v = 1 s) and trial timeline: the abrupt auditory stimulus is delivered at the end of the visual expansion (time 0). Acclimation, 6 min; inter-trial interval, 40 s. Ovals refer to control (grey), auditory (blue), visual (red), multisensory (magenta) randomized trials. (C) Stimulus grid: three visual contrasts (Vmin, Vmed, Vmax), three auditory intensities (Amin, Amed, Amax), their nine combinations (1–9) and a blank condition (Control, zero contrast and zero amplitude). Multisensory stimuli are numbered for reference in G. MC, Michelson Contrast. (D) Eight representative head-velocity traces of motor events; fast events are those exceeding 6 cm/s. Traces are aligned to event start (vertical dotted line); scale, 20 cm/s. (E) Escape response probability for each unisensory stimulus and for each visual–auditory combination (N = 40; Mixed-Effects Binomial GLM, fish identity as random effect, auditory effect 0.91, p < 0.0001; visual effect 0.46, p < 0.0001). (F) Multisensory integration coefficient (MI) for each combination; colour indicates MI. The mean MI across combinations is greater than 0 (Wilcoxon signed-rank test, p = 0.037) and is inversely related to auditory intensity (Gaussian GLM, auditory effect −0.115, p = 0.009), with no significant effect of visual intensity (−0.006, p = 0.90). (G) MI of each multisensory condition as a function of its maximum unisensory response probability; each point is one combination (numbered as in C), averaged across fish. Black line, linear fit with confidence interval; the correlation is negative (slope = −0.50, p = 0.013, R² = 0.61). Shading marks multisensory enhancement (MI > 0) and suppression (MI < 0).

The probability of eliciting a fast escape response (events where head velocity was higher than 6 cm/s, Fig. 1D, short and long latency C-starts pooled, see Methods) increased with the intensity of both visual and auditory stimuli (Fig. 1E). To quantify the effect of combining sensory modalities, we calculated a multisensory integration coefficient (MI) for each of the nine audiovisual conditions. The MI compares the escape probability during multisensory stimulation to the most effective unisensory stimulus (see methods). This analysis revealed that combined audiovisual stimuli significantly increased escape probabilities beyond the maximum unisensory response (Fig. 1F). Interestingly, the magnitude of this enhancement was inversely related to the intensity of the auditory stimulus while the intensity of the visual stimulus had no significant impact on the MI. This was further supported by the negative correlation between the MI of each audiovisual condition and its maximum unisensory response probability (Fig. 1G).

These findings agree with the principle of inverse effectiveness, where weaker unisensory stimuli produce proportionally greater multisensory enhancements^45,46^. For instance, the weakest auditory stimulus (60 dB), elicited responses in 15% of trials while the lowest contrast visual stimulus (Michelson Contrast 0.13) elicited responses in 10% of the trials. When these two weak stimuli were combined (stim. 7 in Fig. 1C), the response probability rose to 33% (Fig. 1E), yielding a mean multisensory integration coefficient of 0.37 (Fig. 1F-G). In contrast, responses evoked by the loudest auditory stimulus (75 dB) did not show multisensory enhancement, showing almost no change compared to its unisensory response probability (Stims. 3, 6 and 9 in Fig. 1G).

Overall, these results confirm that larval zebrafish can combine audiovisual information to decide whether to execute a fast escape response^3^ and that the strongest multisensory enhancement is produced by exposure to the weakest stimuli. However, the mechanistic and neuronal bases of this multisensory integration are unknown. To this aim, we next decided to study the neuronal correlates in the Tectum.

### Auditory Stimulation Elicits Neural Responses in the Tectum

Larval zebrafish are capable of integrating multisensory information during fast escapes (Fig. 1). While visual processing in the fish Tectum has been described, accounts of auditory coding are scarce and focused on the representation of pure tones^27–29,47^. We thus investigated audiovisual representation within the Tectum using a brief and abrupt sound which can evoke fear responses (Fig. 1) to ask whether this classically visual brain region also responds to auditory danger stimuli.

To answer this question, we performed *in vivo* confocal calcium imaging in agarose embedded larval zebrafish at 4-7 dpf expressing the cytoplasmic calcium indicator GCaMP6f (Fig. 2A). We imaged the PVL and the tectal neuropil during the presentation of auditory stimuli and quantified the tectal response during a trial as the proportion of active ROIs (ΔF/F > 1.2 SD above the mean, see methods) during a 2.8 s window after sound presentation (Supp. Fig. 2).

**Figure 2.**
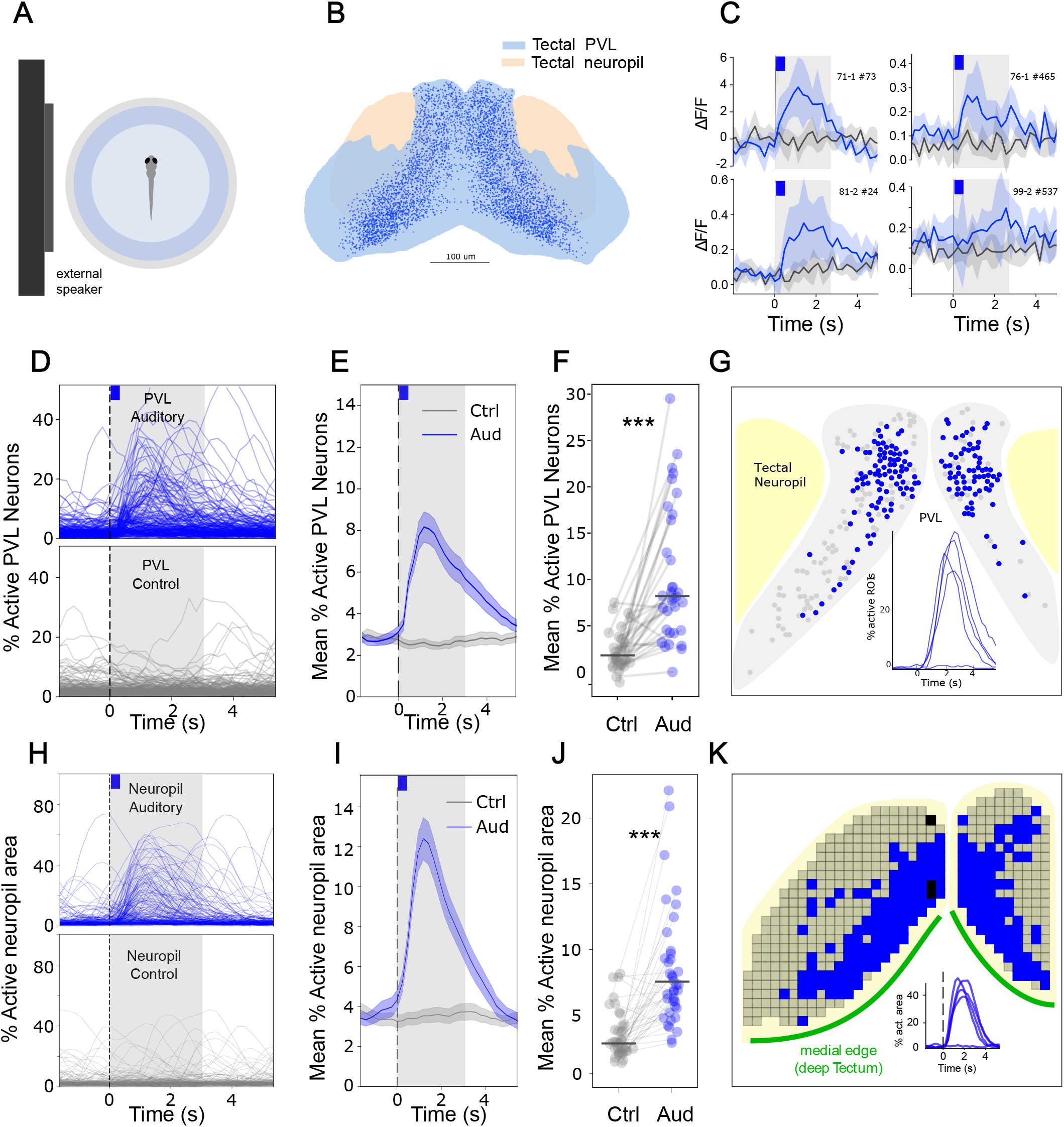
Auditory stimulation elicits neural responses in the Optic Tectum of zebrafish. (A) Configuration for presenting auditory stimuli to agarose-embedded larvae. (B) Anatomical distribution of auditory-responsive PVL neurons (blue dots; 4586 neurons responsive at least once) throughout the PVL of both hemitecta; tectal PVL (blue) and neuropil (orange) are indicated. Scale bar, 100 µm. (C) Auditory responses of four representative PVL neurons (mean ± SE of 5 trials) from four animals; auditory (blue) and control (grey) trials. Vertical dashed line, sound onset; blue bar, stimulus. (D) Upper: percentage of active PVL neurons during auditory trials, one line per trial (5 trials per plane, n_planes = 33). Lower: control (blank) trials. Vertical dashed line, sound onset. (E) Mean (± SE) proportion of active PVL ROIs over time for auditory (blue) and control (grey) trials across experiments; responses are quantified over a 2.8-s window (shaded) after sound onset. (F) Mean proportion of active PVL ROIs during that window per experiment (connected points, 5-trial averages). Ctrl = 2.7%, Aud = 8.9%; Wilcoxon signed-rank test, ***p < 0.001. (G) Distribution of active PVL ROIs during the window for a representative example (auditory, blue; control, black); inset, % active ROIs per trial. PVL outlined in grey, neuropil in yellow. (H) Auditory stimuli increase the proportion of active tectal neuropil area (upper) relative to spontaneous activity (lower) (5 trials per animal, n_planes = 33). (I) Mean (± SE) proportion of active neuropil area over time for auditory (blue) and control (grey) trials across experiments. (J) Proportion of active neuropil area during the window per experiment (connected points, 5-trial averages). Ctrl = 2.9%, Aud = 7.3%; Wilcoxon signed-rank test, ***p < 0.001. (K) Distribution of active neuropil area during the window for a representative example (auditory, blue squares; control, black squares); green line marks the medial edge of each hemi-neuropil (deeper tectal layers); inset, % active area per trial.

We found that the presentation of an auditory stimulus produced a robust activation in the PVL (Fig. 2B) with 4586 neurons responding at least once to the auditory stimulus. Although variable across planes, auditory responsive neurons were distributed along the antero-posterior axis although there was a slight higher density towards the posterior end of the PVL (Fig. 2B). Four examples of individual auditory responsive neurons (mean of 5 trials) from four different animals are shown in Figure 2C. Across 33 imaged planes that met the auditory-response inclusion criterion and were included in the analysis (13851 ROIs, 24 larvae), the mean proportion of active ROIs activated by auditory stimulation showed a sharp increase after onset of stimulation (Fig. 2D, upper panel) while control trials showed a steady and low proportion of spontaneously active ROIs (Fig. 2D, lower panel). Figure 2E shows the time-average of the 33 planes shown in Figs. 2D and indicates a mean simultaneous activation of around 8% of PVL neurons, a proportion notably higher than the basal 3% observed in control trials. Within each plane, the mean proportion of active neurons for auditory trials was also significantly higher than for control trials (Fig. 2F).

The auditory-evoked neural representation in the PVL displays a consistent pattern across trials of the same animal. Figure 2G shows a representative example of the location of responsive neurons in the PVL during auditory (blue) or control (grey) trials respectively. We next asked if auditory responsive neurons were randomly scattered in the tectum or showed some kind of topographical assembly. To test this, we measured the compactness of the auditory response as the mean nearest-neighbor distance among auditory-responsive ROIs. Taking the ratio of compactness of auditory neurons and a random subset of ROIs reveals that auditory-evoked representations are more compact than expected by chance (median: Aud: 0.12, Ctrl: 0.09; compactness ratio is significantly greater than 1; Wilcoxon signed-rank test, *p* < 0.001). This indicates a non-random clustering of responsive neurons which was, however, different from animal to animal.

The tectal neuropil also showed robust auditory responses. To quantify those responses, we subdivided the neuropil into a grid of neuropil ROIs given the absence of clearly identifiable somas. Neuropil ROIs had approximately the same area as ROIs defined in the PVL, however, contrary to PVL ROIs, neuropil ROIs do not correspond to somata but quantify the fluorescence emitted by the neurites located within. Responses in the tectal neuropil during auditory trials (Fig. 2H, upper panel) were stronger and more synchronous, i.e. reached a higher proportion of simultaneously active ROIs, than during control trials (Fig. 2H, lower panel). The neuropil activity showed dynamics that on average reached 13% of its area simultaneously active for auditory trials, which was significantly higher than the 3% active for control conditions (Fig. 2I). Analysis of individual planes showed the same behavior, with a consistently larger area activated for auditory than for control trials (Fig. 2J). Auditory stimulation produced a bilateral and synchronized activity of neuropil ROIs which clustered towards the medial edge of the tectal neuropil (Fig. 2K, blue) while in control (no sound) trials fewer, scattered grid ROIs were active (Fig. 2K, black). The neuropil’s response to auditory stimulation seemed to be concentrated in deeper (closer to the medial edge) layers (Fig. 2K) while activity in the superficial layers was rarely observed. To characterize the spatial distribution of the neuropil activity, we computed the mean distance of the activated neuropil ROIs to the medial edge of the Tectum (green line in Fig. 2K) and found that this distance was significantly smaller than that of a null model created by randomly selecting the same number of ROIs (mean +/- sd: 26.78 ± 10.18 µm vs. 30.56 ± 7.19 µm; Wilcoxon signed-rank test, p = 0.002), indicating a non-random clustering of responsive ROIs, which was consistently observed in most experiments. Beyond clearly showing that there is auditory coding in the Tectum, our results suggest auditory input reaches the Tectum at the deeper layers of the neuropil and contacts the dendrites of neurons whose somata are in the PVL.

In addition, we consider the lack of regionalization of auditory responses along the anteroposterior axis both in the neuropil nor in the PVL suggestive of a distributed auditory input to the Tectum (as opposed to a regionalized area that computes auditory input). In turn this implies that visual input coming for any part of the visual field could be integrated with auditory input.

### Auditory Inputs Enhance Visual Coding in the Tectum

The Tectum participates in the processing of visual looming stimuli^48,49^. Here we investigated whether an auditory cue presented concurrently with a visual looming stimulus (Supp. Fig. 2A) could affect tectal activity (Fig. 3). The visual looming stimulus evoked robust single unit responses in PVL neurons that were visible in the last frames of expansion and reached their peak after the end of the expansion (time 0, Fig. 3A; Supp. Fig. 2C, middle panel). Examples of responses to multisensory stimuli of individual neurons are presented in Figure 3B, showing a similar profile than to visual responses in terms of amplitude and temporal dynamics. Although individual responses were not drastically different, multisensory trials activated a larger number of neurons across experiments compared to visual stimuli alone. Visual trials elicited clear responses that recruited a significant proportion of PVL neurons (Fig. 3C), but multisensory trials induced larger synchronic activation immediately following the auditory stimulus onset (Fig. 3D). Across experiments, the number of active ROIs reached a higher peak in multisensory trials, the proportion of active ROIs in the PVL (shaded area in Fig. 3E) was significantly larger than in visual activation alone trials (Fig. 3F). These results indicate that adding an abrupt auditory stimulus to the end of an visual expansion significantly increases neuronal recruitment in the PVL. Note that as the visual stimulus was delivered from the right side of the animal (Supp. Fig. 2A) it evoked a visual response principally in the contralateral (left) hemitectum (Fig. 3G). Intriguingly, the same visual loom paired with an auditory stimulus recruited a comparable proportion of ROIs in both hemispheres (Fig. 3H). This is consistent, however, with the bilateral activation observed for unisensory auditory responses (Fig. 2).

**Figure 3.**
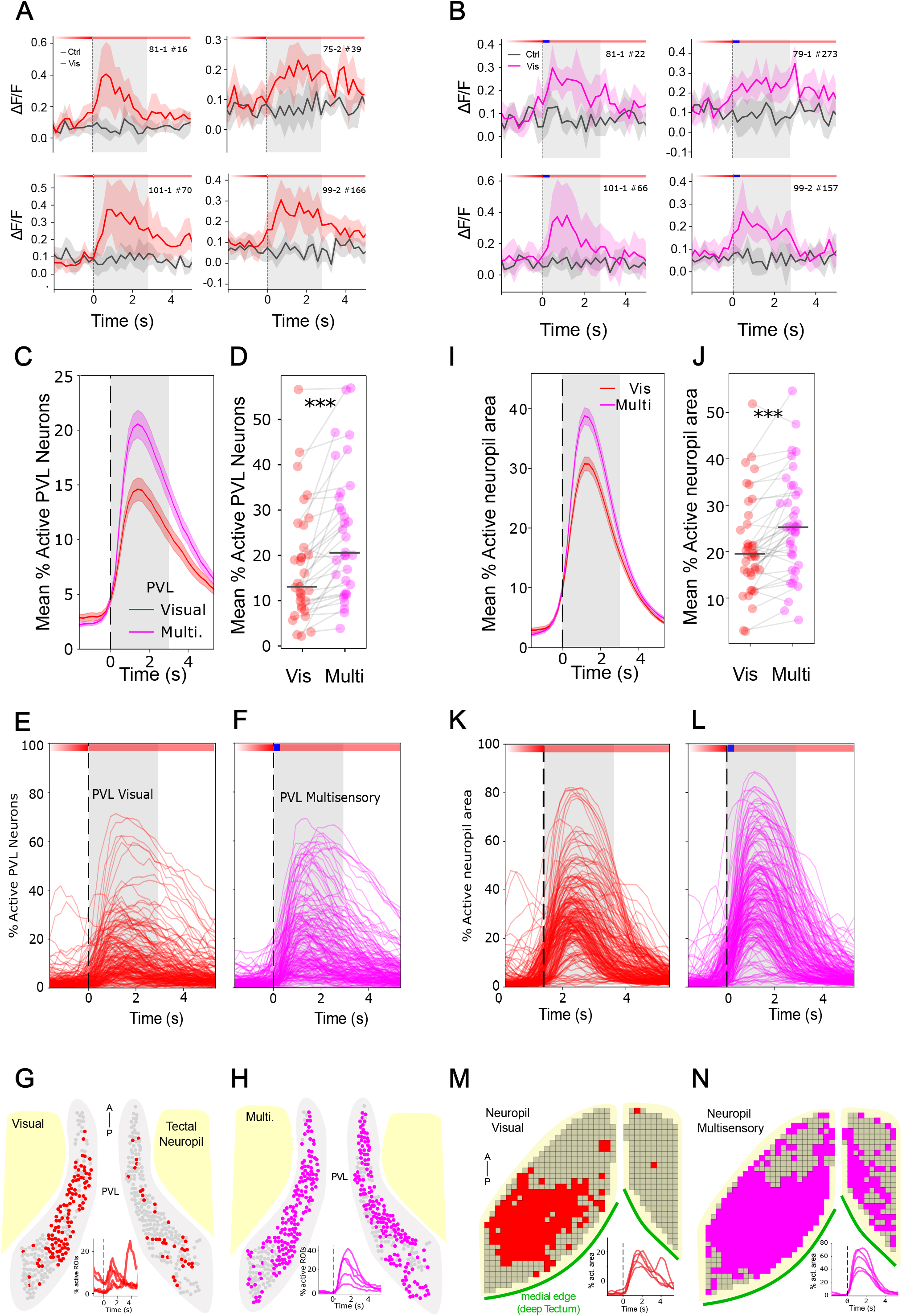
Multisensory stimulation enhances neural responses over a unisensory visual stimulus in the Tectum of zebrafish. (A) Visual responses of four representative PVL neurons (mean ± SD, 5 trials) from four animals; visual (red) and control (grey); Vertical dashed line, last frame of looming expansion. Horizontal red line represents visual loom expansion. (B) Multisensory responses of the same-type neurons; control (grey) and multisensory (magenta). The horizontal line represents visual loom expansion (red) and auditory stimulus (blue). (C) Mean % active PVL neurons during the window, visual vs multisensory, paired per experiment. Vis = 13.0%, Multi = 20.7%; Wilcoxon signed-rank test, ***p < 0.001. (D) Mean (± SE) % active PVL neurons over time, visual (red) and multisensory (magenta). (E) % active PVL neurons for all visual trials, one line per trial (5 trials per plane, n_planes = 33); vertical dashed line, end of loom expansion. Horizontal line, stimulus dynamics. (F) Same for multisensory trials; vertical dashed line, sound onset concurrent with the end of expansion. (G) Distribution of active PVL ROIs during the window for a representative visual example (red dots); inset, % active ROIs per trial; A–P axis. (H) Same experiment for multisensory trials (magenta dots), showing a more distributed and bilateral activation; inset as in G. (I) Mean (± SE) % active neuropil area over time, visual (red) and multisensory (magenta). (J) Mean % active neuropil area during the window, paired per experiment. Vis = 19.3%, Multi = 25.4%; Wilcoxon signed-rank test, ***p < 0.001. (K) % active neuropil area for all visual trials. (L) Same for multisensory trials. (M) Distribution of active neuropil area for a representative visual example (red squares); green line, medial edge (deeper tectal layers); inset, % active area per trial. (N) Same experiment for multisensory trials (magenta squares); activation extends to deeper layers, and the mean distance of active area to the medial edge is smaller than for visual trials (27 ± 8 vs 29 ± 7 µm; Wilcoxon signed-rank test, p = 0.0014).

We next asked if we could also detect multisensory enhancement in the neuropil and if the increased activity observed in the PVL during multisensory trials was matched with an increase in the proportion of active neuropil ROIs. Also, we asked how the anatomical representation of the multisensory responses would relate to the visual representation at input (neuropil) level. Since we observed that adding auditory input to the visual loom produced a more distributed activation across the contralateral PVL and notably of the ipsilateral PVL (Fig. 3H), we wondered if multisensory responses in the neuropil would be also more distributed, suggesting that integration of visual and auditory afferences enhance Tectum activation - at the expense of losing spatial resolution.

We observed a robust increase in the proportion of active neuropil area during multisensory trials compared to visual trials (Fig. 3I-J). The peak proportion of active grid ROIs during multisensory stimulation was 10% higher than in visual only trials (Fig. 3K) while the median proportion of active neuropil area was 6% higher in multisensory trials compared to visual trials (Fig. 3L). Notably, the addition of the auditory stimulus caused the activation of deeper layers of the visible part of the ipsilateral neuropil (ROIs closer to the medial edge, green curves, Fig. 3N), just like we observed with the unisensory auditory stimulus (Fig. 2K). Indeed, the mean distance of the activated neuropil regions to the medial edge was lower in multisensory trials compared to visual trials (Fig. 3M-N, visual, mean +/- sd: 29 +/- 7 um vs. multisensory: 27 +/- 8 um; Wilcoxon signed-rank test, *p* = 0.0014). These results imply that the neuropil’s response to audiovisual stimulation is a combination of visually-evoked activity in superficial and medial layers (Fig. 3M) and auditory-evoked activity in deeper layers (Fig. 2K), serving as the multisensory inputs subsequently processed by the PVL (Fig. 3N). It also confirms that the combination of auditory and visual input not only enhances activity in the hemitectum contralateral to the visual input but produces a significant activation of the ipsilateral neuropil.

### Multisensory Enhancement of Tectal Response Shows Inverse Effectiveness with Stimulus Salience

Numerous studies have reported that the multisensory enhancement of neural responses tends to be inversely correlated to the effectiveness of the inputs when received separately^1,3,11,50^ and thus, particularly effective when individual signals are weak or ambiguous. To ask if multisensory integration in the Tectum also shows inverse effectiveness, we presented the auditory cue (A_max_, Fig. 1C) in combination with a low-contrast visual loom (V_min_. Fig. 4A) or a high contrast loom (V_max_, Fig. 4C) and tested if decreasing the intensity of the visual stimulus increased multisensory enhancement in the zebrafish Tectum.

**Figure 4.**
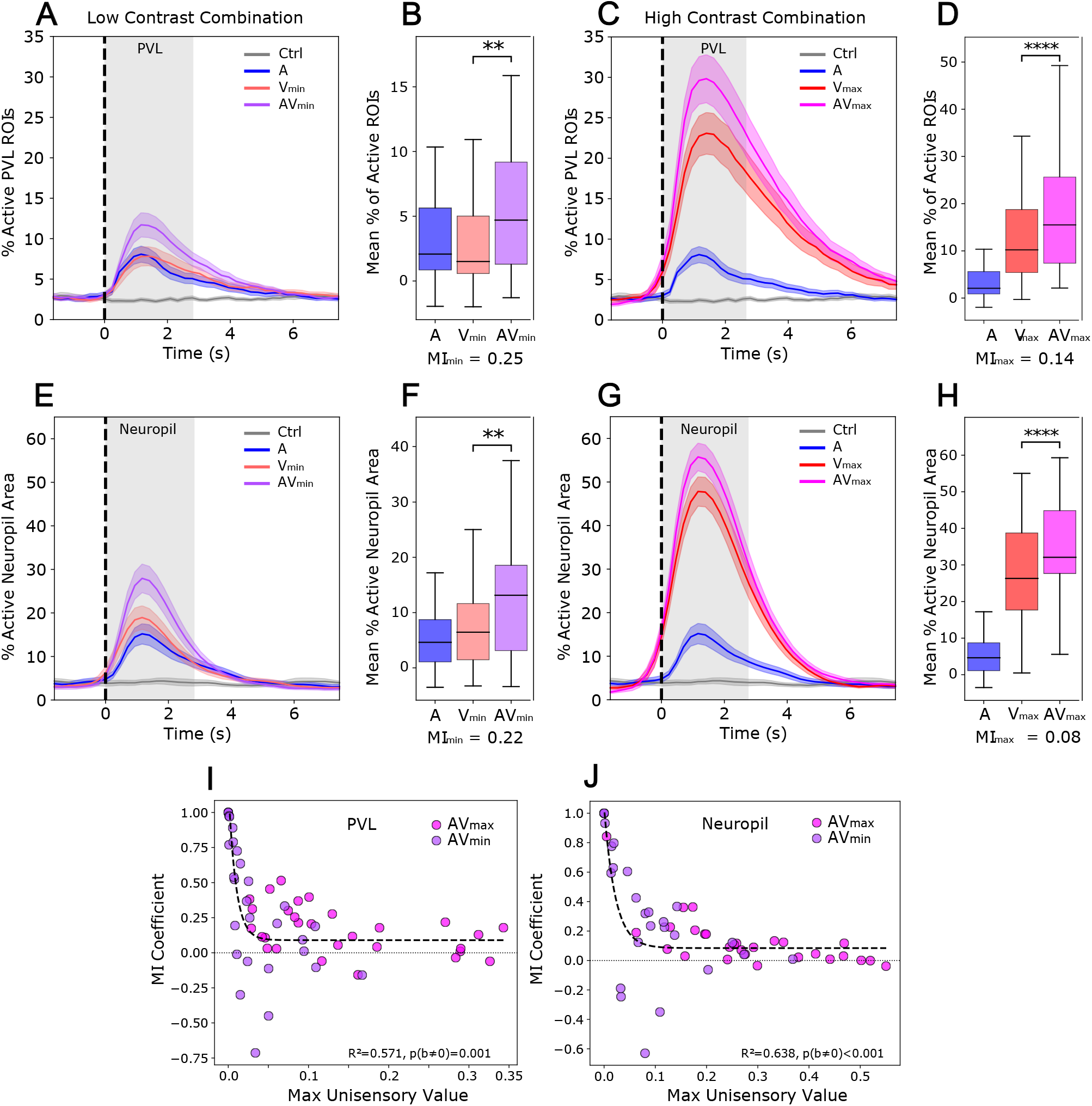
Inverse effectiveness in the zebrafish Tectum. (A) Mean (± SE) proportion of active PVL ROIs over time for the low-contrast combination: control (grey), auditory (A, blue), weak visual (Vmin, light red) and multisensory (AVmin, purple); shaded, analysis window. (B) Mean % active ROIs during the window (control-subtracted) for A, Vmin and AVmin. **p = 0.004 (Vmin vs AVmin); MImin = 0.25. (C) Same as A for the high-contrast combination: control, A, strong visual (Vmax, red) and multisensory (AVmax, magenta). (D) Mean % active ROIs during the window for A, Vmax and AVmax. ****p < 0.0001 (Vmax vs AVmax); MImax = 0.14. (E–F) As A–B for the neuropil; **p (Vmin vs AVmin); MImin = 0.22. (G–H) As C–D for the neuropil; ****p < 0.0001 (Vmax vs AVmax); MImax = 0.08. (I) MI in the PVL vs maximum unisensory activation, per experiment, for the low-contrast (AVmin, purple) and high-contrast (AVmax, magenta) stimulus. Dashed line, single exponential-decay fit (b > 0, p = 0.001, R² = 0.571, n = 58). (J) Same for the neuropil (b > 0, p < 0.001, R² = 0.638, n = 54).

While low-contrast visual stimulus (V_min_) evoked a tectal response in 1.5% of PVL neurons (Fig. 4B), V_max_ recruited 10.2 of PVL neurons (Fig. 4D), a significantly higher proportion (V_min_ vs. V_max_, Wilcoxon signed-rank test, p < 0.001). However, we observed an increase in the mean percentage of active ROIs both with V_min_ combined with an auditory stimulus (Fig. 4B) and with V_max_ combined with an auditory stimulus (Fig. 4D). The MI (Multisensory Index, see Methods) was MI_min_= 0.25 for the low-contrast multisensory stimulus and MI_max_= 0.14 for the high-contrast multisensory stimulus. This larger multisensory enhancement for the low-contrast condition than for the high-contrast condition demonstrates that in the tectal PVL the multisensory integration shows inverse effectiveness.

In the neuropil we observed a similar relationship between visual contrast and the proportion of active neuropil area (Fig. 4E and G) or the mean percentage of active neuropil area (Figs. 4F and H, V_min_= 6.5% vs. V_max_= 26.3%). The effect of combining a visual and an auditory stimulus was stronger for V_min_ (Fig. 4F; MI_min_ = 0.22) but there was also a significant increase for the high-contrast condition (Fig. 4H; MI_max_ = 0.08).

The effective salience of a visual stimulus is not only a function of its contrast, but also depends on gaze direction, attentional state and other factors that can vary between animals and trials. We thus analyzed, for each individual experiment (imaged tectal plane) the relationship between the MI and activity during the visual unisensory condition in the PVL (Fig. 4I) and in the neuropil (Fig. 4J). The inverse effectiveness hypothesis predicts that in experiments where the visual stimulus evokes a weaker response there will be a stronger multisensory enhancement. Indeed, MI and the maximal unisensory response are inversely related for both conditions using a single exponential-decay model, both in the PVL (Fig. 4I) and in the neuropil (Fig. 4J). Low-contrast stimuli are usually perceived as less salient and therefore tend to produce higher MI values compared to the high-contrast condition (purple vs. magenta dots in Figs. 4I and J), although this difference is not significant in the PVL (Wilcoxon signed-rank test, paired by experiment, p = 0.20), possibly due to variability across animals. In the neuropil, however, MI values for the low-contrast condition are significantly higher than for the high-contrast condition (Wilcoxon signed-rank test, paired by experiment, *p* = 0.019). Overall, this shows that the Tectum exhibits stronger multisensory enhancement when the unisensory components of the multimodal stimulus have lower salience, a hallmark of multisensory integration.

### Multisensory Stimulation Recruits Unresponsive PVL Neurons and Boosts Response Probability of Audiovisual Neurons

While the Tectum shows multisensory enhancement at the population level (Fig. 3) and this effect alone could have important consequences on downstream circuits, actual multisensory integration in tectal neurons requires that individual neurons change their response when auditory and visual cues are combined. This is because an increase in the proportion of active neurons could be due to a sum of independent unisensory responses occurring in distinct cells, and not to true multisensory integration at neuronal level. We thus asked if this multisensory enhancement would also be observable at individual neurons, which in our experiments, should be reflected in cells that are only active when auditory and visual inputs are simultaneously present.

We analyzed the response profiles to unisensory and multisensory stimuli of individual PVL neurons (Supp. Fig. 2C). Based only on their response to unisensory stimuli (i.e. excluding multisensory trials) we categorized PVL neurons in four distinct classes: neurons that responded exclusively to auditory stimulation (EA neurons), neurons that responded exclusively to visual stimulation (EV neurons), neurons that responded to both modalities (i.e. to unisensory auditory or unisensory visual stimuli, AV neurons), and neurons that did not respond to either unisensory auditory or unisensory visual stimuli. This last group was further divided into two classes depending on whether neurons were (exclusively) responsive to multisensory stimulation (EM neurons) or unresponsive to all stimulation (U neurons). It’s important to note that, although auditory stimulation produced bilateral activation of the Tectum (Figs. 2G, K), visual stimulation produced preferential activation of the hemitectum contralateral to the screen (Fig. 3G, M). We thus restricted the analysis to the contralateral Tectum (left Tectum, Supp. Fig. 2A-B), where activation of both visual and auditory inputs was robust.

A representative example of this categorization is illustrated in Fig. 5A, where ROIs are color-coded based on the neuron response profile. The spatial distribution and proportions of these neuron types varied across experiments, and we did not find any indication of spatial clustering of the responses (other than retinotopy for stimuli with a visual component, Supp. Fig. 2F). On average, 21% of all PVL neurons analyzed (only left hemitectum) were unresponsive to all stimuli (Fig. 5B, U). On the other hand, 29% of PVL neurons were responsive to visual looms (EV) but never to auditory stimuli while 19% showed the opposite, being only responsive to auditory stimuli (EA). To confirm that these neurons were truly insensitive to their non-preferred modality in our stimulation conditions, we compared their activity during its preferred-modality stimulation with their activity during multisensory trials. Neither EA nor EV neurons showed an increased response to multisensory stimulation (EA, Wilcoxon signed-rank test, *p* = 0.12; EV, Wilcoxon signed-rank test, *p* = 0.89, not shown). This could be explained by the nature of our stimuli and/or the spatial configuration of our setup not being effective (e.g. retinotopic effects for the visual stimulus and possible dependence on sound features for the auditory stimulus), but we cannot rule out that some PVL neurons truly receiving only one kind of sensory input.

**Figure 5.**
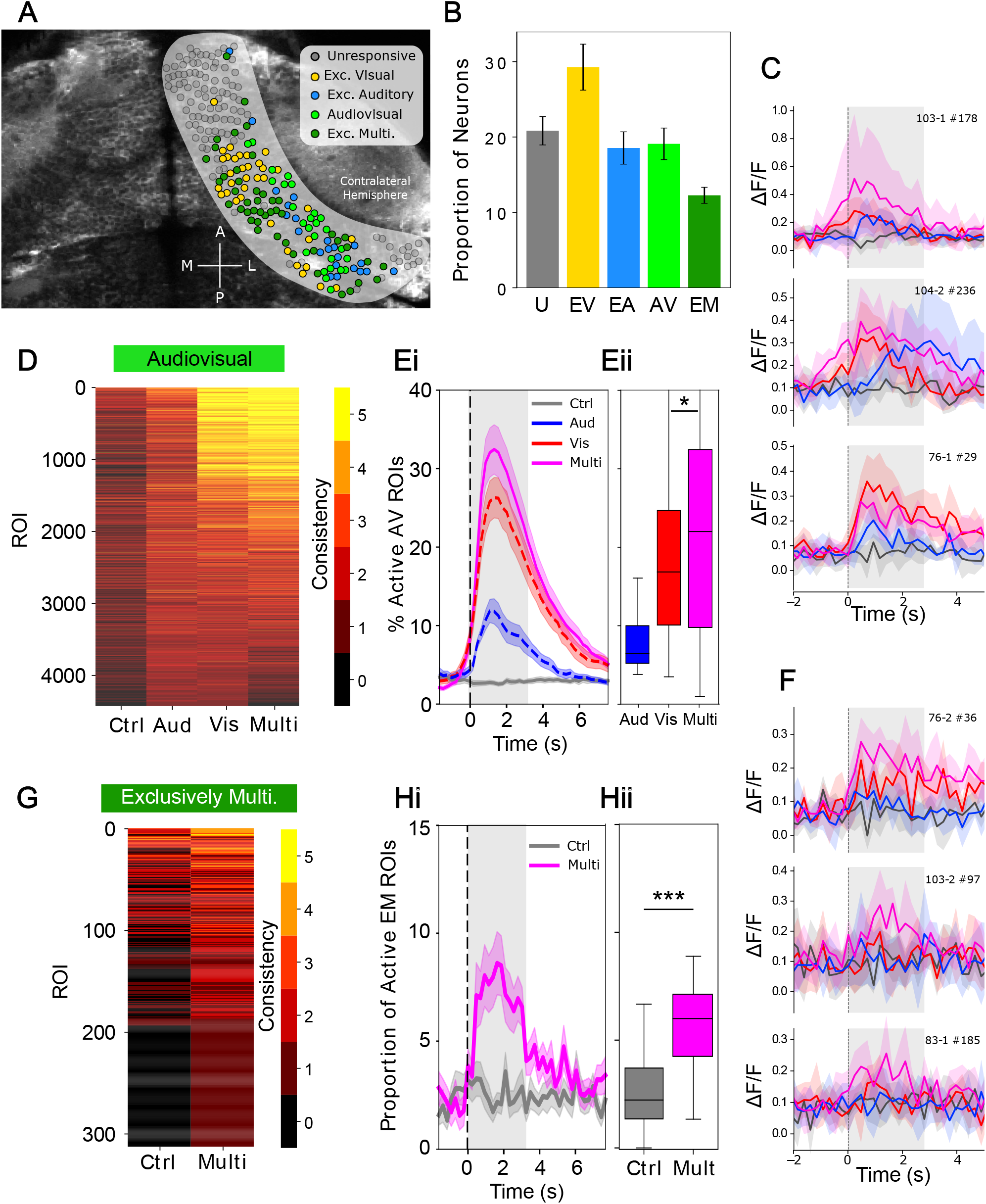
Multisensory stimulation recruits unresponsive PVL neurons and boosts response probability of audiovisual neurons. (A) Representative example of PVL neurons in the hemitectum contralateral to the visual stimulus, colour-coded by response class (Unresponsive, U; Exclusively Visual, EV; Exclusively Auditory, EA; Audiovisual, AV; Exclusively Multisensory, EM), overlaid on the anatomical template; M–L and A–P axes. (B) Proportion of neurons per class across experiments (mean ± SD: U 21 ± 2, EV 29 ± 3, EA 19 ± 2, AV 19 ± 2, EM 12 ± 1). (C) ΔF/F traces of three representative AV neurons for control (grey), auditory (blue), visual (red) and multisensory (magenta) trials, showing multisensory responses that can exceed, match or fall below the visual response. (D) Response consistency of all AV ROIs for control, auditory, visual and multisensory conditions, sorted by mean consistency; AV neurons increase their consistency for multisensory vs. visual trials (Mixed-Effects Linear Regression, fish identity as random effect, effect size 0.15, p = 0.04). (Ei) Mean (± SE) proportion of active AV ROIs over time per condition. (Eii) Mean peak proportion of active AV ROIs for auditory, visual and multisensory trials (Wilcoxon signed-rank test, *p = 0.012). (F) ΔF/F traces of three representative EM neurons per condition (colours as in C), responsive only when both cues are combined. (G) Response consistency of all EM ROIs for control and multisensory conditions, sorted by mean consistency. (Hi) Mean (± SE) proportion of active EM ROIs over time. (Hii) Mean peak proportion of active EM ROIs for control vs multisensory trials (Wilcoxon signed-rank test,***p < 0.001).

On average 19% of PVL neurons responded to the visual loom but also to unisensory auditory trials (Fig. 5B, AV), a proportion much larger than the <1% previously reported for non-danger audiovisual cues^27^. We asked if these AV neurons that are activated by both unisensory auditory and unisensory visual stimuli would increase their responsiveness when presented with those stimuli simultaneously, or if they would show audiovisual suppression as previously shown for non-danger cues^27^. Example traces of three individual AV in Figure 5C show that responses to multisensory stimuli could be significantly larger, similar or smaller than to unisensory visual stimuli. To systematically quantify AV responses to auditory, visual and multisensory stimuli, we calculated the consistency of each AV neuron as the number of trials in which it was active for a particular stimulus. This value represented the number of trials of a given stimulus condition in which a neuron was active, ranging from 0 (for a consistently inactive ROI) to 5 (for a ROI that was activated on each of the 5 trials for a particular condition). We calculated the consistency of each AV neuron individually for each condition: control (no stimulation), auditory, visual and multisensory (Fig. 5D). Given that visual responses were stronger and more consistent than auditory responses across our experiments, we compared the multisensory consistency against the visual consistency. AV neurons increased their consistency (i.e. their individual probability of response) for multisensory vs. visual trials. This is also apparent when calculating the proportion of active audiovisual ROIs over time (Fig. 5Ei), where the peak multisensory activity of these neurons is significantly higher than their peak activity for either visual (or auditory) unisensory stimuli (Fig. 5Eii). These results parallel those observed for the whole population of PVL neurons (Figs. 3E-F) and critically show that simultaneous auditory and visual input can enhance response probability at the individual neuron level.

Strikingly, we found that about 12% of PVL neurons were silent during all ten unisensory trials but responded to (at least once) exclusively multisensory stimulation (Fig. 5B, EM). To ensure that these responses were not due to spontaneous activity, we compared the activity of these neurons during multisensory trials against their activity during unisensory or control trials (Fig. 5F). Based on their categorization, auditory and visual responses of EM neurons were indistinguishable from control trials, therefore, to quantify EM response we computed their consistency in control and multisensory trials. EM neurons were more consistently active during multisensory stimulation (Fig. 5G) and their peak response probability was as well significantly higher than when compared to control trials (Fig. 5H). This shows that multisensory stimulation can recruit cells which, although receive inputs from both modalities, are silent unless those inputs are activated simultaneously. Note that to be conservative, we restricted the analysis to the contralateral hemitectum, where visual responses are stronger. Had we included the ipsilateral hemitectum, the number of EM neurons (and their relative response to visual stimuli) would be significantly higher (Supp, Fig. 2C-D).

We next asked if the activity of AV neurons would form distinct representations of visual and auditory stimuli (i.e. different responses to auditory and visual unisensory stimuli), and, if so, how multisensory representations would relate to unisensory representations (Supp. Fig. 3). To answer this question, we performed temporal principal component analysis (t-PCA) on the ΔF/F data from individual AV PVL neurons during visual, auditory, and multisensory trials within a 1.8 second window after stimulation time (which encompasses the period where populational responses reaches their peak, Fig. 5Ei). This allowed us to transform N-dimensional activation patterns (where N is the number of PVL neurons in a particular experiment) into trajectories in a low-dimensional space which encode information about the temporal dynamic of the tectal representation for each trial.

Two representative examples from two different animals are shown in Supp. Fig. 3A-B. Plots of the first two components of the PCA show trajectories for trials that showed a significant response, i.e. there can be up to 5 trajectories for each type of stimulus (lines correspond to PVL trajectories in response to visual, auditory and multisensory stimuli). All trajectories were normalized to start at the origin, which represents the Tectum state before stimulation. Trajectories corresponding to visual and auditory trials diverged in PCA space (median angle between visual and auditory trajectories was 60°), suggesting that the tectal representation to visual and auditory stimuli are different.

To describe the variability of the multisensory patterns of activity and compare it to their unisensory counterparts in all experiments, we calculated the median unisensory direction for each unisensory stimulus Supp. Fig. 3A-B) and evaluated the angle of each multisensory trajectory with both unisensory vectors (Supp. Fig. 3C). Most multisensory patterns of activity align with either the visual or the auditory representation, i.e. each pattern of activity has a small angle with the visual vector and a larger one with the auditory vector or vice versa, with the visual mode being more common (73% visual, 27% auditory). To quantify the separation between unisensory representations, we treated each complete trajectory (spanning 8 post-stimulus frames) as a single vector in a 16-dimensional space (2 coordinates x 8 time points). We then computed the average Euclidean distance between all pairs of visual-auditory trajectories (inter-modality distance) and compared it to the average distances within the same modality (intra-modality distances, i.e., visual-visual and auditory-auditory). The inter-modality distance was significantly greater than intra-modality distances (Supp. Fig. 3D) confirming that visual and auditory stimuli evoke distinct and separable population activity patterns in the PVL.

To determine whether the multisensory response aligned with one of the unisensory inputs or had a distinct activity pattern, we compared the multisensory trajectories against their unisensory counterparts. First, we computed the average Euclidean distance between multisensory and auditory trajectories (inter-modality) and compared it to their respective intra-modality distances. The multisensory-auditory distance was significantly greater than both the intra-multisensory and intra-auditory distances (Supp. Fig. 3E). Conversely, when we performed the same comparison between multisensory and visual trajectories, the inter-modality distance was not significantly different from the intra-modality distances (Supp. Fig. 3F). This analysis confirms that the population response to the multisensory stimulus is closer to the visual-only response, as opposed to the auditory-only response.

### Multisensory Integration in the Tectum Is Associated with Enhanced Motor Responses in the Hindbrain

We have shown that the zebrafish Tectum integrates visual and auditory information, enhancing neural responses. Does this multisensory enhancement have a functional behavioral value? The Tectum is known to project to premotor regions in the hindbrain, both in zebrafish and in other vertebrate species^25,26^. Some hindbrain neurons receive input from the Tectum to evoke fear-related responses, such as (but not restricted to) the Mauthner cell-initiated C-start responses^22,51,52^. It is therefore possible that audiovisual integration in the Tectum could affect premotor neurons and, consequently, influence the execution of behavioral responses. For this reason, we next explored the relationship between the enhancement of tectal activity during multisensory trials and behavioral output.

Although larvae were agarose-embedded to restrict brain movement during image acquisition they still exhibited small, sudden movements that were registered during the recording. These sudden movements in the x-y plane are routinely corrected using a motion correction algorithm before ROI extraction^53^. Interestingly, we noticed that these motion events were more frequent during stimulus presentation (Fig. 6A). Frame-by-frame displacements of confocal images larger than 4 SD above the mean displacement of each experiment (see methods) were considered to be motion events (Fig. 6B). Across all imaged planes (n=33) we found that these motion events were more common within the period following visual (4%) or auditory (6%) stimulation than during spontaneous activity (1%). Crucially, the probability of eliciting motion events during multisensory trials (12%) was significantly higher than the maximum unisensory (auditory) condition (Fig. 6A). These results parallel the multisensory enhancement we observed for escape responses in behavioral experiments (Fig. 1E-F), although response probabilities here are much lower, likely because of the differences in the conditions of stimulus administration.

**Figure 6.**
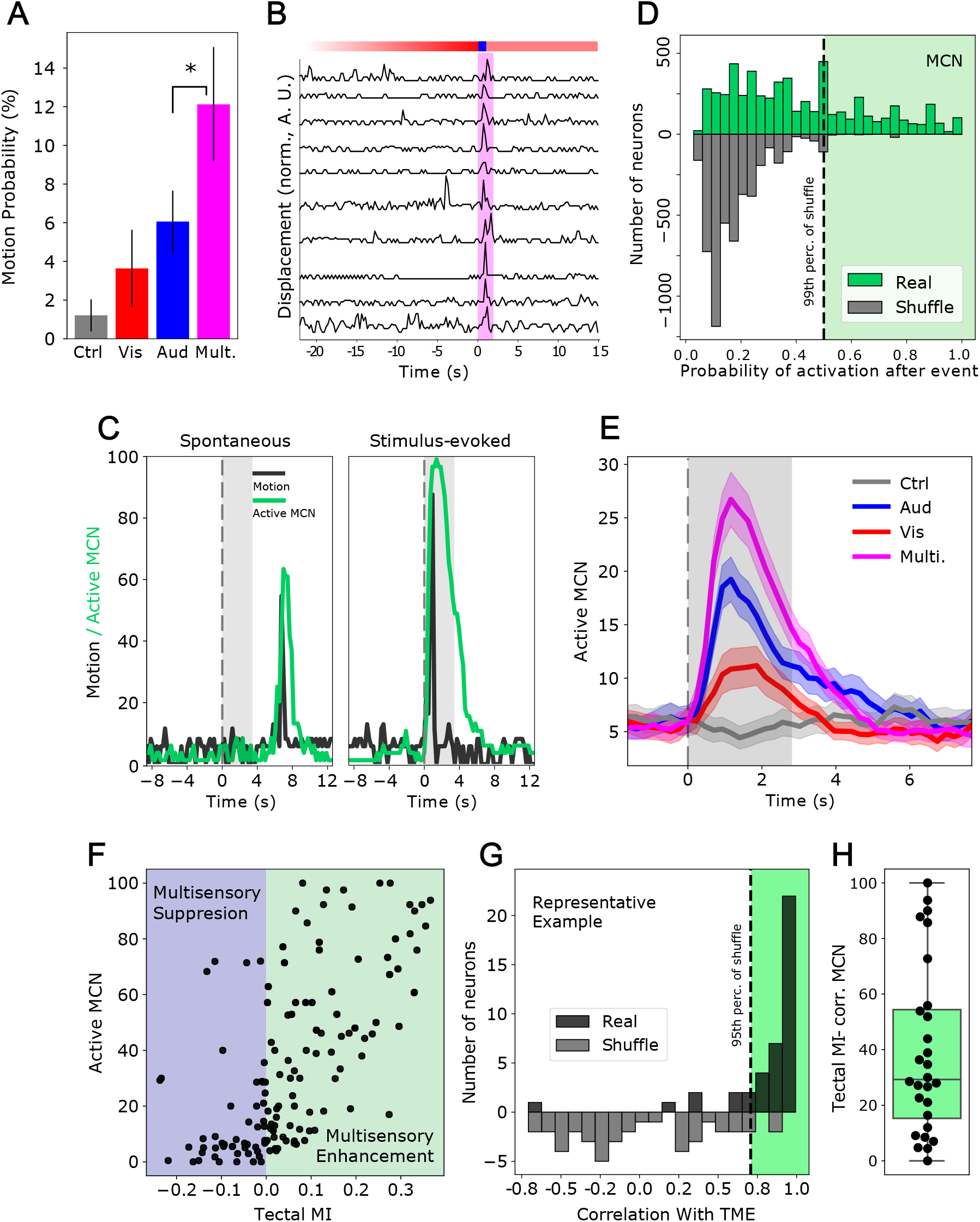
Motion-correlated neurons in the Hindbrain are mostly responsive to multisensory stimulation when tectal activity shows multisensory enhancement. (A) Motion-event probability for each stimulation condition; bars are means across experiments (paired t-test, Aud vs. Multi, t = 2.26, *p = 0.029). (B) Ten representative frame-by-frame image-displacement traces (motion-correction output) in trials where a motion event was detected around stimulation (pink band); a motion event is a displacement > 4 SD above the experiment mean. (C) Synchronous activation of hindbrain motion-correlated neurons (MCNs) tracks motion events. Left: representative spontaneous event (outside the stimulation window); black, image displacement (a.u.); green, proportion of active MCNs. Right: representative stimulus-evoked event. (D) Distribution of the per-neuron activation probability after a motion event, for all recorded hindbrain neurons across experiments, for real events (green) vs shuffled event times (grey); only events outside the stimulation window were used. Vertical dashed line, 99th percentile of the shuffle. (E) Mean proportion of active MCNs over time across experiments for control, auditory, visual and multisensory stimuli; the multisensory response exceeds the auditory one (Mann-Whitney U test, p = 0.037). (F) Active MCNs as a function of tectal MI; each dot is a trial. Stronger tectal multisensory enhancement (MI > 0) is associated with a higher proportion of active MCNs (linear regression on MI > 0, coefficient = 0.65, p < 0.001), whereas suppressive tectal trials (MI < 0) are not. (G) A subset of MCNs is associated with tectal MI: distribution of the correlation between each MCN’s mean ΔF/F and tectal MI across the 5 multisensory trials in a representative experiment; real (black) vs shuffled trial order (grey). Vertical dashed line, 95th percentile of the shuffle; green shading, MCNs above it. (H) Proportion of MCNs whose ΔF/F is significantly correlated with tectal MI, per experiment (each point one experiment).

If these events represent actual motor responses this should be reflected in activity of hindbrain neurons. We initially focused on neurons primarily located in rhombomeres 1 and 2, which could be imaged simultaneously with tectal activity. More caudal rhombomeres were outside the field of view. To identify candidate premotor neurons, we first calculated the activation probability of each hindbrain neuron considering only spontaneous motion events (i.e. events occurred in the absence of sensory stimulation, Fig. 6C, left panel). We next produced a shuffled distribution of activations for each trial and each neuron, where motion event times were randomized to eliminate real correlations and compared the real and shuffled distributions. We defined motion-correlated neurons (MCN) as those neurons with an activation probability higher than 99th percentile of the shuffled distribution (Fig. 6D).

We confirmed that this population of hindbrain neurons was active during both spontaneous and stimulus-evoked motion events (Fig. 6C). Notably, the proportion of MCNs that became active in response to sensory stimulation was higher for auditory than for visual trials and showed multisensory enhancement over the auditory condition (Fig. 6E), reproducing the pattern observed in the motion events (Fig. 6A).

The enhancement in the proportion of active MCN of rhombomeres 1 and 2 of the hindbrain after multisensory stimulation (Fig. 6E) was reminiscent of the multisensory enhancement we observed in the Tectum (Figs. 3E, 4B). We thus asked if the level of activity of the MCNs correlated with MI observed in the Tectum. For each multisensory trial, we calculated tectal MI, similarly to what we did in Fig. 4B to quantify multisensory enhancement (see methods). We observed that when tectal MI was zero or negative—indicating no enhancement or suppression—the activation of MCN in the hindbrain was weak and not significantly associated with MI (Fig. 6F, linear regression, *p* = 0.716). However, when tectal MI was positive, there was a strong positive association between MI and MCN activity in the visible rhombomeres (Fig. 6F). This shows that there is a positive relationship between multisensory enhancement in the Tectum and hindbrain activity linked to behavioral output. Crucially, this relationship is rectified as a function of integration valence: only positive multisensory effects in the Tectum are associated with an increased probability of MCN activation in the anterior hindbrain.

To determine if this association between tectal MI and hindbrain activity was driven by all or by a subset of these MCNs, we calculated the mean ΔF/F of the five multisensory trials for each MCN and computed the correlation coefficient between its mean ΔF/F and the tectal MI value for the corresponding trials (Tectal MI - corr. coef. MCN). This metric indicates, for each experiment, how strongly each MCN was correlated with tectal MI during multisensory stimulation. In Fig. 6G black bars represent the real distribution of Tectal MI - corr. coef. MCN for an example experiment while gray bars show the results when shuffling trial order to remove all significant associations. All neurons with coefficients larger than the 95th percentile of the shuffled distribution (green shaded area in Fig. 6G) were considered to be correlated with tectal activity. Across experiments, the proportion of MCNs exceeding this threshold varied (Fig. 6H) but on average 30% (+/- 22% SD) of MCNs were positively correlated with tectal multisensory integration. The correlation between tectal multisensory integration and activity in the anterior hindbrain neurons driving motor responses provides not only a functional link between these two structures but a mechanistic basis for the multisensory enhancement observed at the behavioral level (Fig. 1). The Tectum neurons receive visual and auditory afferences at different depths on the neuropil (Figs. 2, 3). On neurons insensitive to unisensory stimulation, multisensory input could result in suprathreshold responses, resulting in a larger number of active PVL neurons. In turn, this increased tectal input to premotor neurons such as the Mauthner cell could increase the probability of a motor response, effectively lowering the threshold for threat detection.

However, motor circuits responsible for fast escape responses driven by the M-cell network are distributed along the hindbrain^54^. While the M-cell series, located in rhombomeres 4, 5 and 6^55^, initiate the escape response, the dynamics of the following escape swim depends on activation of downstream networks that include cranial relay neurons (located caudally, in rhombomeres 7 and 8) which in turn project to forward swimming networks located in the midbrain^56^.

Although simultaneous recordings from the tectum and rhombomeres caudal to rhombomere 2 was not possible in our setup (and thus we cannot directly correlate tectal activity with midcaudal MCNs), we still decided to extend the characterization of hindbrain activity beyond rhombomere 2 to have a more comprehensive picture of how multisensory stimuli drive activity in areas directly involved in motor behaviors. For this we recorded rhombomeres 1 to 7 (Supp Fig. 4A) in 30 planes from 18 larvae and identified MCNs as before (Fig. 6). We analyzed only planes with enough motion events for reliable MCN extraction (16 planes from 11 larvae). When considering all identified MCNs we obtained a similar pattern of motion probability to unisensory and multisensory stimuli (Supp. Fig. 4B) than in the anterior hindbrain (Fig. 6A). Multisensory stimuli recruited a higher proportion of MCNs than auditory or visual stimuli, while response probabilities to all stimuli were higher. The proportion of MCNs in each rhombomere (Supp. Fig. 4C) varied between 30 and 70%, as well as the response probability to each type of stimuli (Supp. Fig. 4D). In all rhombomeres with the exception of rhombomere 5, the multisensory stimulus had the highest response probability. This yielded positive MI (Supp. Fig. 4E), notably, MCNs of rhombomere 1 (whose activity was correlated with the level of MI in the tectum, Fig. 6F) show the highest MI. Caudal rhombomeres 6 and 7 also showed positive MI. Unfortunately, the M-cell series does not express the calcium sensor in the transgenic fish used in this study and thus we cannot directly assess their response.

## DISCUSSION

This study provides the first comprehensive evidence of multisensory enhancement elicited by audiovisual stimuli across neural and behavioral levels in zebrafish. Using visual and auditory danger cues, we observed robust multisensory integration in the Tectum. We identified motion-correlated neurons in the hindbrain which also show an increased response to multisensory stimulation. On a behavioral level, multisensory stimulation significantly enhanced escape response probability. This paradigm enables an examination of how multisensory processing at the neural level influences overt behavioral decisions that, to date, has not been explored in zebrafish and remains relatively rare in the literature on other species.

### Auditory, visual and multisensory units in the zebrafish Tectum

Work in zebrafish and other species suggests that deeper tectal layers receive non-visual sensory inputs^14,15,28,57^, supporting the role of the Tectum as a multisensory integrative structure. Our findings build on this concept and expand previous observations of Thompson et al. (2016)^27^, demonstrating that auditory information reaches the deeper layers of the zebrafish tectal neuropil (Figure 2K). Response within these layers is robust and enhanced during multisensory stimulation (Fig. 3L-N), suggesting that integratory circuits that have been found in other vertebrates are anatomically conserved in zebrafish.

Here we used brief abrupt auditory stimuli that triggers escape responses in freely moving larvae. Our stimulus configuration was functionally non-directional, probably sensed as a vibrational stimulus (Supp. Fig. 1 C-G) which produced bilateral activation of the deeper layers of the tectal neuropil (Fig. 2K). It is interesting to note that although, as expected, visual stimuli produced stronger activation on the contralateral hemitectum (Fig. 3G-M), when combined with the auditory stimulus this spatial asymmetry was lost (Fig. 3H-N) was significantly reduced. This suggests that integration of auditory input activates the tectum independently of visual input. Behaviorally, this would mean that animals are able to detect a potential threat with a higher sensitivity (i.e. before, or even if it is weak) at the expense of source location accuracy, which would reduce directionality of the escape. Alternatively, direction could be computed downstream the tectum, before the motor command is triggered. Further experiments are necessary to test this hypothesis.

We also identified a significant presence of neurons in the Tectum that respond to multiple sensory modalities, a finding documented in other species but scarcely observed in zebrafish. Previous reports using non-threatening cues estimated that less than 1% of tectal cells were responsive to both visual and auditory stimulation^27^. Here we found a much larger proportion of audiovisual-responsive cells which averaged 19% across experiments (Fig. 5B-C). These audiovisual neurons were responsive to unisensory visual stimuli, but exhibited increased activation probability under multisensory conditions (Fig. 3D), suggesting that neurons that receive audiovisual inputs have a higher firing probability if those signals arrive simultaneously. We also found a proportion of tectal neurons that were only recruited when the auditory stimulus was simultaneous with the end of the visual expansion and never responded to either visual or auditory stimuli independently (EM neurons, Fig. 5B, F-H). It is important to note that these neurons are likely not exclusively multisensory for all possible visual and auditory stimuli (i.e. stronger unisensory stimulation might evoke responses in these units), but under our stimulation regime they only show suprathreshold activity when both inputs are present.

Previous reports on the zebrafish Tectum^27^ identified a suppression of visual responses presented simultaneously with auditory stimuli. In contrast, we did not find suppression but enhancement of visual responses when the auditory component was present (Fig. 4 and 5). One possible explanation for this discrepancy is the nature of the stimuli used in each study. Unlike Thompson et al., which used long pure tones intended to cover the larval hearing range, our study employed abrupt auditory stimuli specifically associated with aversive contexts that evoke robust escape reactions (Fig. 1). Our rationale was that the more naturalistic our stimulus configuration was, the clearer the valence of the stimulus was going to be and thus more effective to reveal any integration mechanisms. We used a stimulus configuration widely used to mimic a predator attack, i.e. a dark image expanding on a light background that couples with a strong abrupt noise at the end of the expansion (Fig. 1B). In nature, an aerial predator - a bird which will look dark against the sky-that dives into the water - making a brief but strong splash as it breaks the water surface to catch a fish would constitute such a stimulus configuration. The Tectum is known to differentiate between cues of varying ecological significance, such as prey and predator signals^34,35,58^ specifically enhancing integration for aversive or appetitive stimuli, while neutral or mismatched cues may lead to multisensory suppression. Indeed, we found that when multisensory integration in the Tectum was suppressive, activity in MCN was minimal (Fig. 6F) suggesting that no motor response would be evoked. Further studies are needed to investigate the effect of stimulus valence in tectal multisensory integration.

About 20% of tectal neurons did not respond to any of our stimuli (Fig. 5B). This is unsurprising given the topographic organization of the Tectum and the spatially localized nature of our visual stimulus within the animal’s visual field. However, previous studies in zebrafish indicate that certain neurons respond exclusively to non-audiovisual modalities^27^. Specifically in zebrafish, tectal neurons known to be selectively responsive to water flow stimuli receive inputs from the lateral line nerve^59^. Thus, lack of response in a subpopulation of neurons might be indicative of a modality not tested in our study.

Finally, it is noteworthy that we observed a pronounced inverse effectiveness pattern, particularly in the neuropil (Fig. 4E-F), where the dendrites of tectal neurons are located^16,25^. This anatomical arrangement suggests that subthreshold effects of dendritic integration could be most visible in this area compared to the somas in the PVL. The presence of inverse effectiveness in the zebrafish Tectum mirrors findings in the superior colliculus of mammals and the Tectum in other vertebrate models^11,45,50^, suggesting an evolutionary conserved multisensory integration mechanism.

### A functional relationship between tectal integration and behavior

In this work, we introduce a methodology for extracting motor event data when simultaneously recording tail movements is not feasible. We demonstrated that these movement events are reliably evoked by both visual and auditory stimuli and show multisensory integration consistent with our behavioral findings (Fig. 6A-B). Additionally, we showed that, even with this inferred movement data, it is possible to identify neurons correlated with these motor events, including both spontaneous movements and stimulus-evoked responses (Fig. 6C-D). In freely moving and in agarose embedded fish, auditory stimuli evoked slightly higher response rates than visual stimuli (Fig. 1D and 6A). The fact that motion events of agarose-embedded fish were associated with the presentation of danger stimuli and that they exhibit multisensory enhancement qualitatively similar to freely-swimming fish suggests that these events represent escape-like attempts.

We report for the first time a relationship between the multisensory enhancement observed in the Tectum and the activity of motion-correlated neurons in the hindbrain, which receive projections from the Tectum^25,26,57,60^. Hindbrain neurons drive premotor activity—prepontine neurons in rhombomere 1 are implicated in evoking long latency C-starts^44^ or the Mauthner cells, in rhombomere 4, are responsible for the short latency C-start. Here we show that multisensory enhancement in the Tectum correlates with an increase in the likelihood of premotor activity (Fig. 6-G). This, in turn, may contribute to the behavioral enhancement we observe, both in agarose-embedded and freely moving animals (Fig. 1), implying a causal link between tectal and behavioral multisensory enhancement.

Our work shows that the zebrafish Tectum integrates aversive multisensory stimuli, producing both sublinear and supralinear enhancement at the single-neuron level and at the population level. This multisensory activity is closely related to the activation of functionally identifiable neurons in the hindbrain. The multisensory integration we describe using aversive danger cues contrasts with prior reports using neutral stimuli. The difference can be attributed to the inherently different processes triggered by stimuli of different valence. It could be hypothesized that while neutral stimuli are be filtered out or even produced an inhibitory integration process (multisensory suppression), stimuli with high valence (negative in our study but positive if we had used prey-like stimuli) recruit distinct neural circuits, including additional attentional resources and/or modulatory input from fear-processing areas. It could be that valence of the stimulus acts as a gate for multisensory integration: while neutral or irrelevant stimuli triggers multisensory suppression (or no integration), high valence stimuli triggers multisensory enhancement which functionally implies increasing the gain to the information provided by ecologically relevant stimuli. These results, beyond extending our current understanding of the tectal neural integration capabilities, stresses the critical importance of using naturalistic stimuli to analyze complex brain processes, which might otherwise run unnoticed.

## RESOURCE AVAILABILITY

### Lead Contact

Requests for further information and resources should be directed to and will be fulfilled by the lead contact, Violeta Medan.

### Materials availability

This study did not generate new unique reagents.

### Data and code availability

All data reported in this paper will be shared by the lead contact upon request. All original code has been deposited at Medanś Lab GitHub and is publicly available at [DOI] as of the date of publication. Any additional information required to reanalyze the data reported in this paper is available from the lead contact upon request.

## LIMITATIONS OF THE STUDY

Our results indicate that multisensory stimuli increase activity both in the Tectum and hindbrain and that the proportion of active MCNs increases with the level of multisensory enhancement in the Tectum (Fig. 6F). However, we only have indirect (correlative) evidence of a relationship between MSI in the tectum and level of recruitment of MCNs of rhombomere 1. Multisensory stimuli also could directly project to MCNs in the rest of the rhombomeres (auditory input directly contacts the M-cell in rhombomere 4), and therefore we cannot rule out that increased recruitment of MCNs produced by multisensory stimuli occurs independently of the Tectum integration we report. The direct contribution of auditory integration in the tectum on MCNs activity could be tested expressing an inhibitory opsin in the neuropil to block synaptic transmission when the auditory stimulus reaches the tectum. Changes in activity of hindbrain neurons or motor responses (in tail-free animals) would be indicative of a direct tectal contribution. Also, although in our behavioral analysis we only included fast escape responses (but did not differentiate between short and long latency C-starts^40,44^) it could be possible that some of the MCN events that we included in the correlation analysis of tectal MSI are non-escape struggling events. Indeed, most motor events recorded outside the stimulation window (Fig. 6C) belong to the latter category. Therefore, although our results are robust they are still correlational and should be interpreted with caution.

## ACKNOWLEDGMENTS

This work was supported by PICT 2020-2136 FONCYT-ANPCYT (VM), PIP

11220200102799CO (VM), CONICET, UBACyT 20020190200319BA (VM), UBA. NM was supported by graduate fellowship from CONICET. The authors thank Drs. Szczupak, Berón de Astrada, Belluscio and Otero Coronel and members of the Medan lab for discussion and for reading previous versions of this manuscript. Authors also thank Lirane Moutinho and Ángel Vidal for technical assistance.

## AUTHOR CONTRIBUTIONS

Conceptualization, NM, VM; Methodology, NM, VM; Investigation, NM, VM; Data curation, NM; Software, NM; Validation, NM, VM; Visualization, NM, VM; Funding acquisition, VM; Writing-original draft, NM, VM; Writing-review and editing, NM, VM.

## STAR★METHODS

### KEY RESOURCES TABLE

| REAGENT OR RESOURCE | SOURCE | IDENTIFIER |
| --- | --- | --- |
| Chemicals, peptides, and recombinant proteins |  |  |
| 4% ultrapure low-melting agarose | Sigma-Aldrich | Cat# A9414-25G |
| Deposited data |  |  |
| Code for behavioral analysis | Medan lab | https://github.com/.... |
| Code for imaging analysis | Medan lab | https://github.com/.... |
| Experimental models: Organisms/strains |  |  |
| ZebraFish: Tg(elavl3:GCaMP6f)jf1Tg | <sup>61</sup> | ZFIN: ZDB-ALT-150916-3 |
| ZebraFish: mitfa<sup>b692/b692</sup> | <sup>62</sup> | ZFIN: ZDB-FISH-150901-6335 |
| Software and algorithms |  |  |
| DeepLabCut | <sup>63</sup> | https://github.com/deeplabcut/deeplabcut |
| CalmAn library | <sup>53</sup> | https://github.com/flatironinstitute/CalmAn |

## METHOD DETAILS

### Animal husbandry

We used larval zebrafish (*Danio rerio*) for this study. All animals were group-housed in 5 l aquaria of recirculating water, in a 14-10 h light–dark cycle (lights on at 8 am) in a temperature controlled room at 24-26 °C. All experiments were conducted between 9 am and 5 pm. Embryos were raised in E3 medium (5mM NaCl, 0.17 mM KCl, 0.33 mM CaCl2, 0.33 mM MgSO4) buffered with 10 mM HEPES (pH 7.3) at 28 °C, with medium changes every day. Larvae were fed AP100 (Zeigler Larval AP100 Diet) daily from 7 days post-fertilization (dpf) onward. We utilized larvae aged 4 to 7 dpf of AB background carrying the mitfa−/− mutation and Tg(elavl3:GCaMP6f)jf1Tg^64^ for calcium imaging experiments. For behavioral experiments, we tested 80 larvae between 4 and 15 dpf. The behavioral analyses presented in this study were restricted to larvae aged 4–7 dpf (n = 40; 10 larvae per age), matching the developmental range used for calcium imaging. For calcium imaging experiments, a total of 68 larvae, aged 4–7 dpf, underwent tectal imaging. An additional 18 larvae underwent extended hindbrain imaging. Across all behavioral and imaging experiments, 166 larvae were tested or imaged; 75 larvae contributed data to the analyses reported here. All experimental procedures were conducted in compliance with current regulations and were approved by the Institutional Animal Care and Use Committee of the Exact and Natural Sciences School, University of Buenos Aires (protocol 152).

### Behavioral Experiments Setup

Behavioral assays were conducted in an arena consisting of a translucent plastic weighing boat of flat bottom, measuring 40 x 40 x 7 mm (Fig. 1A). The arena was filled with buffered E3 medium and was transilluminated using an infrared LED grid (55 x 55 mm, 36 LED units) and passed through opal diffusion glass to achieve more uniform irradiance. An overhead high-speed monochrome camera (720 x 540 px, 437 fps, Blackfly BFS-U3-04S2M-CS, FLIR) equipped with a 35 mm objective (59872, Edmund Optics) recorded larval behavior. The behavioral setup was mounted on an anti-vibration table and enclosed with opaque curtains to avoid unintended stimulation.

### Calcium Imaging Experiments Setup

For calcium imaging experiments zebrafish larvae were immobilized by complete embedding in 4% ultrapure low-melting agarose (A9414-25G, Sigma) and submersed in E3 medium. Calcium imaging was performed using a Zeiss LSM-900 confocal microscope equipped with a 20× water-immersion objective lens (Zeiss W N-ACHROPLAN 420957) driven by a 488 nm laser. Emission was collected through a 509 nm dichroic mirror and 490-620 nm emission filter using a GaASP-PMT detector. For each larva, we acquired a z-stack encompassing the whole volume of both Tecta and adjacent structures (317 x 317 x 300 μm^3^, 101 planes, interplane interval of 3 μm) to align brain images with the Z-Brain Atlas (Randlett et al., 2015). In each larva, at least one coronal plane of the Tectum (317 x 317 μm) was recorded at 4.3 Hz to capture the neural activity in response to a complete set (see below) of sensory stimulation. We also obtained a higher-resolution reference image of each plane used for ROI segmentation. In some experiments, we performed two sequential acquisitions of coronal planes of the Tectum at depths located between 30-60% of the total dorsoventral extension of the Tectum obtaining a total of 114 planes of 68 animals. If an acquired plane suffered substantial drift or gross motion artifacts it was excluded from the analysis.

### Stimulus Presentation

#### Visual Stimuli

Visual stimuli were delivered using a 1.5-inch OLED display module (Waveshare) connected to an Arduino Due microcontroller. The display presented a looming stimulus (Adafruit_SSD1327 and custom C++ code) consisting of a dark circle that expanded over a white background. Looming stimuli are widely used to mimic the approaching of an object in a collision course or the attack of a predator and are effective to trigger escape responses^65^. Their efficacy depends on the contrast, velocity and apparent size of the stimulus^35,39,66^. In our study, the looming stimulus had an l/v ratio of 1, where l is the radius of the virtual object and v is the approach speed^66^. The expansion phase lasted 25 s, followed by a 10-s period where the circle remained at its maximum size. In all analysis, time 0 s signals the end of the visual expansion (Fig. 1B).

As multisensory enhancement depends on stimulus salience (Martorell & Medan, 2022), we designed stimuli of varying intensity. To change the intensity of the visual stimulus, we adjusted the contrast of the expanding circle against a constant white background. The contrast is expressed as:

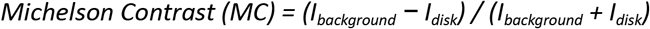

where I_background_ is the irradiance of the background (33 mW/m²) and I_disk_ is the irradiance of the dark disk. The irradiance was measured using a Tektronix J17 photometer equipped with a J1812 sensor. Based on preliminary experiments we defined three levels of visual contrast (Fig. 1C): V_min_ (MC = 0.13), V_med_ (MC = 0.29), V_max_ (MC = 0.59) which showed progressive increase in the probability of eliciting an escape response (Supp. Fig. 1E).

#### Auditory Stimuli

Auditory stimuli were produced by a speaker (Thonet & Vander HK096-03451) placed on foam insulation padding and positioned 10 cm from the arena (Fig. 1A and Supp. Fig. 1A). The intensity of the auditory stimuli was measured on air using a sound level meter (Yu Fung Model YF-20) and reported in decibels (dB re. 20 μPa).

The speaker emitted a single-cycle sine wave at 220 Hz generated and transmitted directly from the sound card of the stimulation computer. The sound frequency was selected based on previous results by us and others showing robust acoustic responses (both behaviorally and in octaval nuclei) within the 100-400 Hz range^47,67,68^. To modulate the intensity of the auditory stimuli, we varied the amplitude of the sine wave, defining three amplitudes: A_min_ (60 dB), A_med_ (69 dB), A_max_ (75 dB). When tested behaviorally (N = 40), these stimuli had respectively low, medium and high salience in terms of the probability of eliciting an escape response (Supp. Fig. 1B). Visual and auditory stimuli were triggered and synchronized using a custom script written in Processing (Processing Foundation).

#### Multisensory Stimuli

We combined the 3 visual and 3 auditory unisensory stimuli to create 9 multisensory stimuli (Fig. 1C). For multisensory stimuli, the beginning of the single-cycle auditory stimulus was timed to coincide with the end of the visual looming expansion, simulating the simultaneous arrival of visual and auditory cues indicative of an approaching threat (Fig. 1B). Previous work from our lab showed that this stimulus configuration effectively evokes escape responses in fish^3^. Visual and auditory stimuli were presented from sources located at 90° (behavioral experiments, Fig. 1A) or 135° (calcium imaging experiments, Supp. Fig. 2A) from each other. Analysis of behavioral responses to auditory (Supp. Fig. 1C-D) and visual (Supp. Fig. 1E-F) unisensory stimuli showed no strong directional component in the trajectory (first 50 ms after response onset, Supp. Fig. 1C and F) nor in the angle of maximum curvature after response onset (Supp. Fig. 1D and G).

### Experimental Procedures

Prior to the start of experiments, larvae were given at least 6 minutes to acclimate to the experimental setup. In behavioral experiments individual larvae were exposed to each of the uni- or multisensory stimuli once, and an additional control condition with no stimulation in random order totaling 16 trials with an intertrial interval (ITI) of 40 s (Fig. 1B-C). In calcium imaging experiments, larvae were exposed to 5 repetitions of a subset of these conditions: V_max_, A_max_, their multisensory combination and the control condition (random order, 20 trials in total, ITI = 40 s). Some calcium imaging experiments (Fig. 4) also included the V_min_ in combination with A_max_ (30 trials in total, ITI = 40 s).

### Behavioral Data Analysis

#### Pose Estimation Using DeepLabCut and Behavioral Event Classification

Larval motor behavior was tracked using DeepLabCut^63^. We trained a ResNet-50 convolutional neural network where the output layer was replaced with a pose estimation layer designed to predict the location of 12 anatomical landmarks as scoremap grids: one point for each eye, two points for the swim bladder, and eight points along the tail. Training was performed on a single NVIDIA GeForce RTX 3060 GPU with 12 GB of VRAM.

An initial training dataset was created by annotating 152 frames from a library of 1,280 behavioral recordings. Frames were chosen to maximize variability in body poses, positions within the arena, fish sizes, and lighting conditions. For a second training iteration, we annotated an additional 125 frames where the model’s initial predictions had low likelihood scores (high uncertainty in model predictions).

The annotated dataset was split into training and validation sets, with 95% of the images used for training and the remaining 5% reserved for model evaluation. Using pose estimation data we calculated 36 kinematic parameters, such as linear head velocity or total body curvature (Supp. Table 1). Discrete activity events, or bouts, were identified by detecting peaks in the linear head velocity time series. Events exceeding a threshold of 6 cm/s were classified as fast responses, as visual inspection of video recordings showed that all events exceeding this threshold were C-starts. (Fig. 1D). We confirmed that this threshold was a robust indicator of a fast escape by kernel PCA analysis^69^. For all events exceeding the 6 cm/s threshold, we estimated 36 kinematic parameters (Supp. Table 1) based on DLC trajectories. Kernel PCA analysis using these 36 parameters allowed us to define two clusters that corresponded to short (SLC) and long (LLC) latency C-starts^40^. We confirmed that the clusters indeed corresponded to SLC and LLC computing maximum angular velocity and stage 1 duration for each event of each cluster that did not differ from those reported by Burgess and Granato (2007)^40^. However, we combined all fast escapes for analysis purposes in Figure 1 since both types of fast escapes showed comparable results (not shown).

#### Quantification of Multisensory Integration

In both behavioral and imaging experiments, multisensory integration was quantified by comparing the activity elicited during multisensory stimulation with the activity observed during the most effective unisensory stimulus^45^. For behavioral experiments, activity was measured as the probability of eliciting a fast response after time 0 (onset of the auditory stimulus or the last frame of the mooming expansion, Fig. 1B). In imaging experiments, it was measured as the proportion of active ROIs in a 2.8 s window after time 0.

The multisensory integration coefficient (MI) was calculated using the formula:

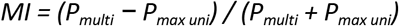

where P_multi_ represents the activity during the multisensory stimulus, and P_max uni_ represents the activity during the unisensory stimulus that elicited the largest response. An MI value of zero indicates no effect of multisensory integration beyond the maximum unisensory response. A negative MI value indicates multisensory suppression, whereas a positive MI value indicates multisensory enhancement.

### Calcium Imaging Data Processing and Analysis

#### Image Registration, Motion Correction and Cell Segmentation

Calcium imaging recordings were processed to correct for motion artifacts and registered to the higher-quality template using the CaImAn library^53^. We next used custom Python code for semi-automated Region Of Interest (ROI) detection based on the anatomical features of the images. Note that the calcium sensor expression is cytosolic and therefore cell nuclei are not fluorescent. The Tectum is organized into the PVL, a deep layer containing most neuronal somas, and the neuropil, a superficial layer where dendrites and afferent axons converge. We detected ROIs in three specific brain regions: the tectal neuropil, the periventricular layer of the Tectum (PVL), and the hindbrain (Supp. Fig. 2B). To estimate the proportion of the visible neuropil area exhibiting significant activity we analyzed the neuropil both as an entire undivided region and by subdividing it into a grid 5 μm x 5 μm to capture regional changes in neurite activity.

#### Fluorescence Normalization, Time Series Analysis and Inclusion Criteria

For each ROI we calculated fluorescence time series by averaging the pixel intensities within the ROI for each recorded frame. These time series were then normalized to ΔF/F, defined as:

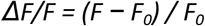

where F is the fluorescence intensity at a given time point, and F_0_ is the baseline fluorescence (calculated as the 8th percentile of a 60-frame (approx. 14 s) window preceding the current frame. An example of all PVL neurons identified in a simple plane from a single experiment are shown responding to auditory, visual and multisensory stimuli in the upper, middle and lower panel of Supp. Fig. 2C. ROIs were binarized and classified as responsive if they exhibited activity greater than 1.2 SD above the mean ΔF/F value in at least one of the five trials of that stimulus during the stimulation window (Supp. Fig. 2D). ROIs inactive during the entire recording session were excluded from subsequent analysis. All 114 planes for the 68 animals had ROIs responding to visual stimuli, however we only included experiments where at least one ROI responded in at least 3 out of 5 auditory trials (yielding 33 planes from 24 animals, Supp. Fig. 2E). The consistency of a ROI’s response was quantified by counting the number of trials in which the ROI was active during the stimulation window (i.e. the consistency values ranged 0 to 5). We adopted this conservative inclusion criteria because in tectal planes where there were no auditory-responsive neurons, comparing unisensory vs. multisensory responses was not possible. Responses were quantified in a 2.8 s (12 frames) window after stimulus presentation (Supp. Fig. 2C-D, magenta shaded area).

To examine population-level neural representations, we performed temporal principal component analysis (t-PCA) on ΔF/F time series from PVL neurons (Supp. Fig. 3). The analysis was restricted to a window 1.86 s following stimulus onset, which encompassed the peak of population responses. This dimensionality reduction allowed us to project n-dimensional neural activity patterns (where n is the number of PVL neurons recorded in each experiment) onto a two-dimensional space, facilitating the visualization and comparison of population trajectories across sensory conditions.

### Identification of Motion-Correlated Neurons

Although embedded in agarose, larvae showed clear motor activity as displacements of the image (Fig. 4B). The activity often was time-locked to the frames immediately following sensory stimuli. The frame-by-frame displacement data obtained from motion correction analysis were used to construct a continuous estimate of the animal’s movement over time (Fig. 4B). To detect motor events, we calculated the mean displacement for each experiment, averaged over all recorded frames. Any frame where the displacement exceeded 4 SD above the experiment’s mean displacement was considered a motion event. This threshold captured large, abrupt displacements indicative of motor-related activity while minimizing the influence of minor shifts that could result from slight image drift.

### Statistical Analysis

All statistical analyses were conducted using Python (v3.9) with the SciPy and Statsmodels libraries. Statistical significance was assessed at a threshold of *α* = 0.05. Binomial Generalized Linear Models (GLMs) were used to evaluate the effects of visual and auditory stimuli on escape response probabilities. Mixed effect models were used in some cases to factor in the variance accounted for by individual differences. Model significance was tested using likelihood ratio tests, and parameter estimates are reported with their associated p-values. The significance of MI values across conditions was assessed using one-sample t-tests. Wilcoxon Signed-Rank Tests were used to compare paired conditions in imaging experiments (e.g., auditory vs. control, multisensory vs. visual), while non-paired tests were performed using Mann-Whitney U-tests. Z-Tests for Regression Slopes were used to compare slopes between different conditions. All analyses were corrected for multiple comparisons where applicable using the Holm-Bonferroni method. Error bars represent standard error unless otherwise specified. Graphs and visualizations were generated using Matplotlib and Seaborn libraries. The code is open-source and available as stated in the Resources Availability section.

## SUPPLEMENTAL INFORMATION

Document S1. Supplemental methods and Figures S1–S4.

---

**Supplementary Figure 1.**
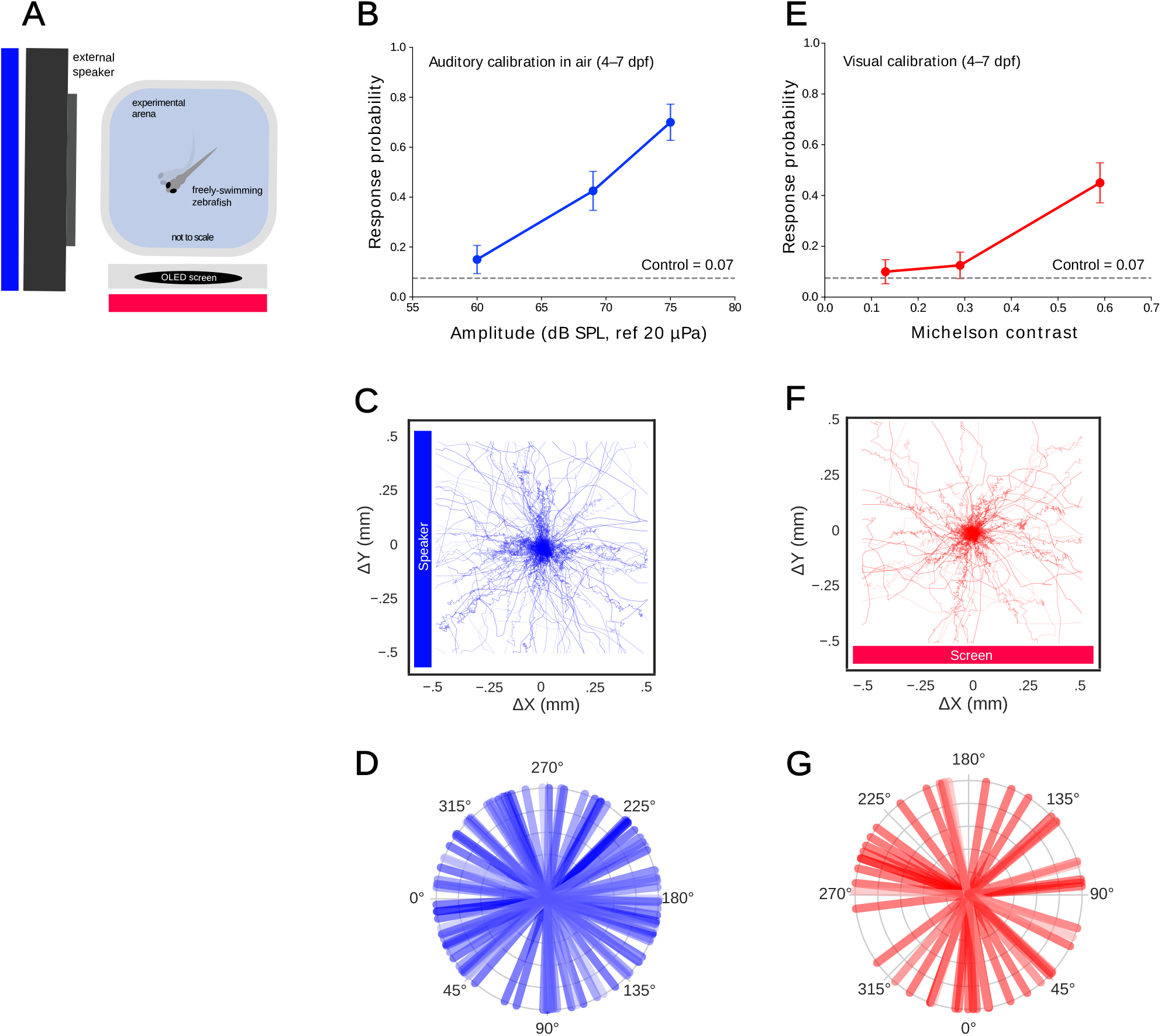
Stimulus calibration and non-directionality of unisensory escape responses in freely-swimming larvae. (A) Behavioural arena (external speaker, experimental arena and OLED screen; not to scale). (B) Auditory calibration in air (4–7 dpf, N = 40): escape response probability vs sound amplitude (60, 69, 75 dB SPL); dashed line, blank control probability (0.07). (C) Auditory-evoked escape trajectories during the first 50 ms after response onset (ΔX–ΔY). (D) Angle of maximum curvature after auditory-evoked responses (polar plot). (E) Visual calibration (4–7 dpf, N = 40): escape response probability vs Michelson contrast (0.13, 0.29, 0.59); dashed line, blank control (0.07). (F) Visual-evoked escape trajectories during the first 50 ms after response onset (ΔX–ΔY). (G) Angle of maximum curvature after visual-evoked responses (polar plot). Neither modality showed a strong directional component in trajectory (C, F) or curvature (D, G).

**Supplementary Figure 2.**
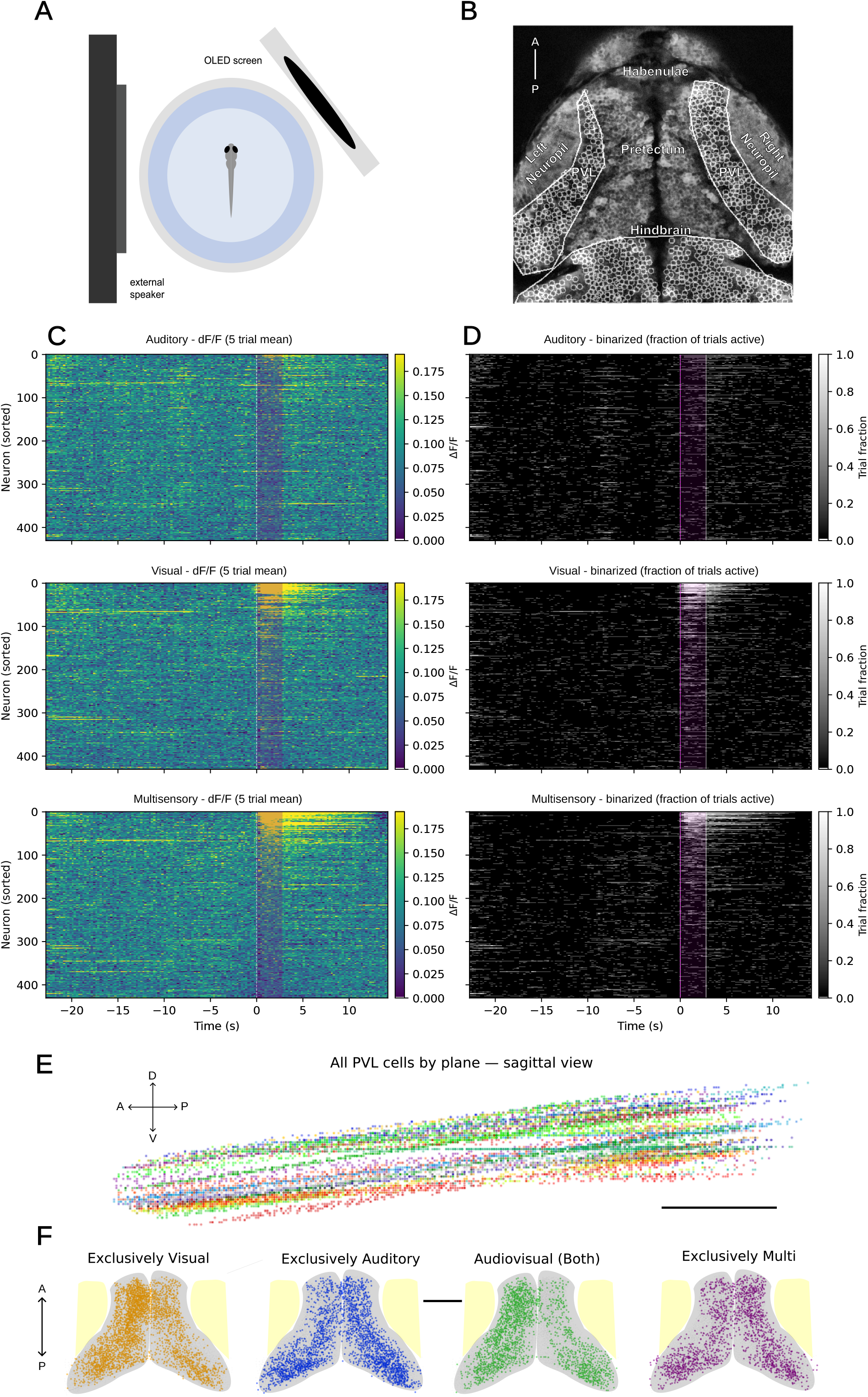
Imaging configuration, ROI segmentation and population responses in the tectal PVL. (A) Imaging configuration: agarose-embedded larva with the external speaker and the OLED screen positioned at 135° from each other, the visual loom delivered from the right side. (B) Representative confocal plane with segmented ROIs and anatomical landmarks (Habenulae, Pretectum, left and right neuropil, PVL, Hindbrain); A–P axis. (C) Trial-averaged ΔF/F (5 trials) of all PVL neurons from one plane, sorted, for auditory (top), visual (middle) and multisensory (bottom) stimuli; time 0, stimulus onset. (D) Same neurons, binarized as the fraction of the 5 trials in which each ROI was active; magenta band, 2.8-s response window. (E) All segmented PVL cells across included planes (sagittal view), coloured by plane; D–V and A–P axes. Scale bar, 50 µm. (F) Spatial distribution of the four response classes (Exclusively Visual, Exclusively Auditory, Audiovisual, Exclusively Multisensory); neuropil in yellow. Scale bar, 100 µm.

**Supplementary Figure 3.**
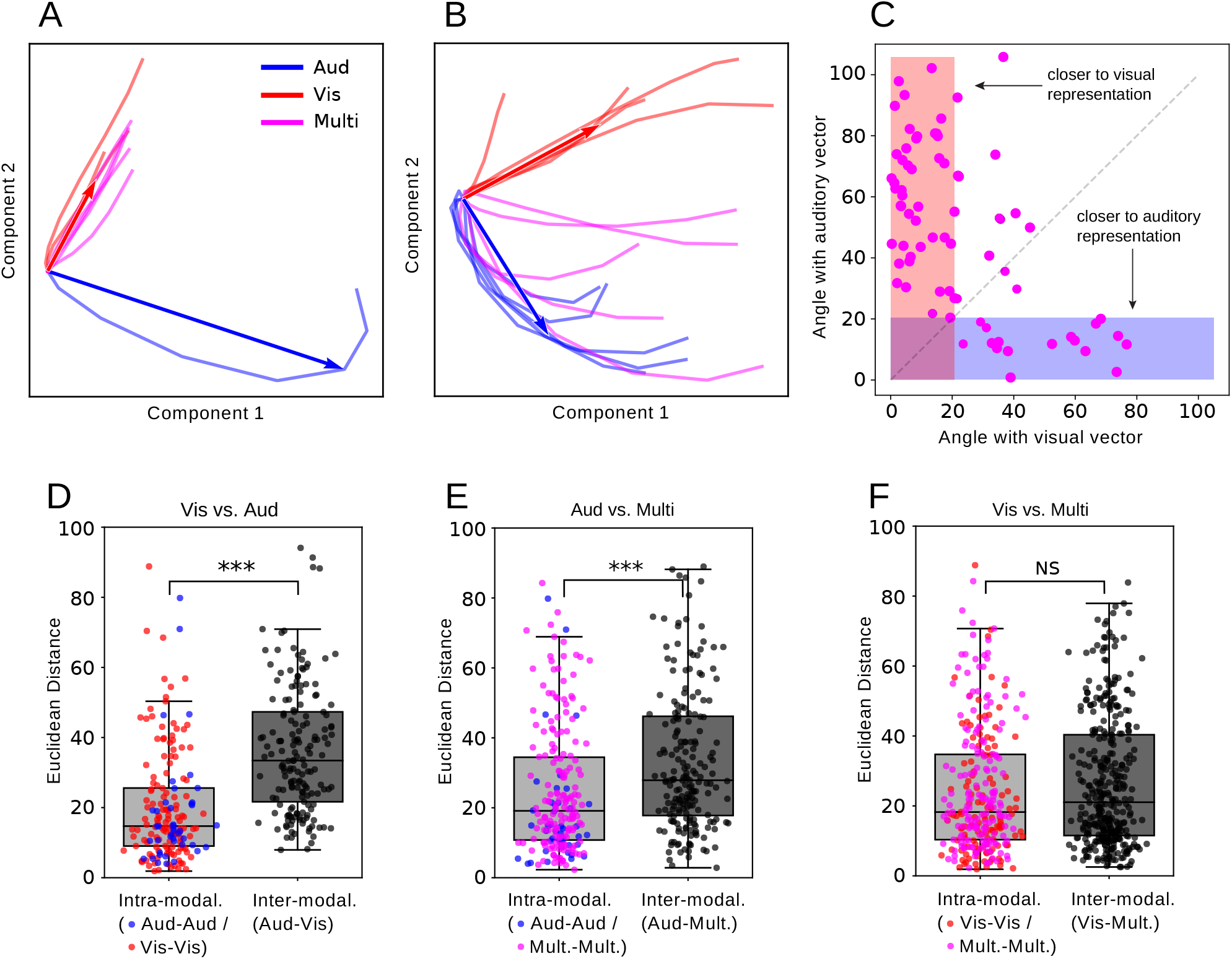
Multisensory representation in the tectal PVL usually aligns closely to a single unisensory representation. (A–B) Representative t-PCA results for two experiments: trajectories of visual (red), auditory (blue) and multisensory (magenta) trials in the space of the first two principal components, over the 1.86 s after stimulus onset. Arrows, median direction of the visual and auditory trajectories. (C) Angle between each multisensory trajectory and the visual (x-axis) and auditory (y-axis) median directions of its experiment; each point is one multisensory trial; grey dashed line, equal distance to both representations; shaded bands mark trajectories closer to the visual or to the auditory representation. (D) Euclidean distance between trajectories of the same unisensory stimulus (Intra-modal: visual–visual, red, or auditory–auditory, blue) and between unisensory stimuli (Inter-modal: auditory–visual, black); dots, distances of each trajectory to all others of the target class within an experiment; boxplots, means of intra- and inter-modality distances (Mann-Whitney U test, ***p < 0.001). (E) As D, between multisensory and auditory trajectories (Mann-Whitney U test, ***p < 0.001). (F) As D, between multisensory and visual trajectories (Mann-Whitney U test, p = 0.12, NS).

**Supplementary Figure 4.**
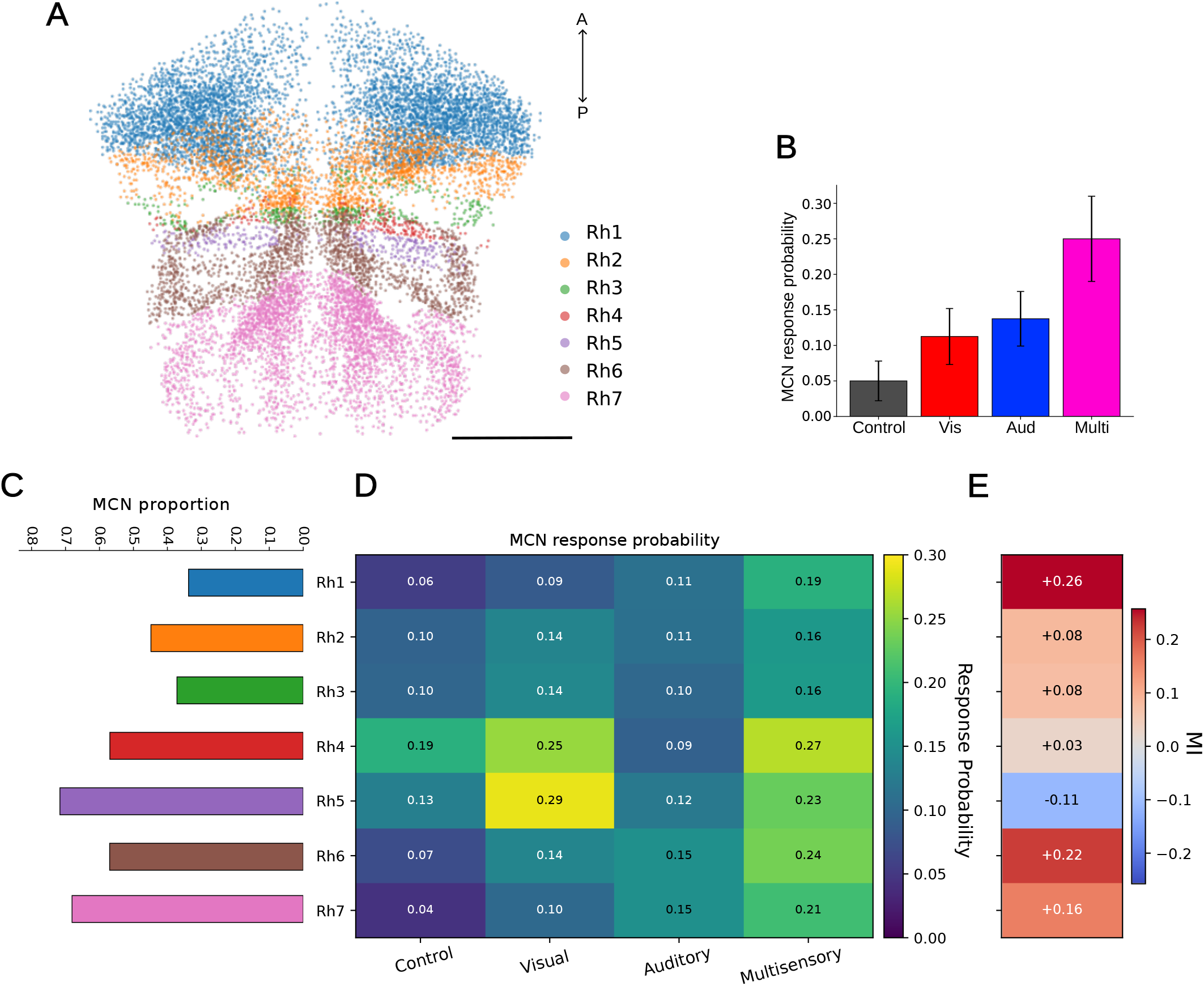
Multisensory recruitment of motion-correlated neurons across hindbrain rhombomeres 1–7. (A) Spatial map of hindbrain MCNs coloured by rhombomere (Rh1, n = 2659; Rh2, n = 970; Rh3, n = 234; Rh4, n = 72; Rh5, n = 127; Rh6, n = 921; Rh7, n = 1996); A–P axis. Scale bar, 100 µm. (B) Mean (± SE) MCN response probability by condition (control, visual, auditory, multisensory), pooling all MCNs (n = 16 planes from 11 fish); the multisensory condition recruits the highest proportion of MCNs. (C) Proportion of MCNs per rhombomere (ranging ∼30–70%). (D) MCN response probability per rhombomere and condition; the multisensory stimulus evokes the highest probability in every rhombomere except Rh5 (where the visual stimulus dominates). (E) Multisensory integration coefficient (MI) per rhombomere; Rh1 shows the highest MI (+0.26), Rh6 (+0.22) and Rh7 (+0.16) are also positive, and Rh5 is the only negative one (−0.11).

**Supplementary Table 1:** Kinematic descriptors of escape behaviors extracted from pose estimation data. Head Velocity: instantaneous speed of the head, calculated as the displacement of the midpoint between both eyes over time. End-Tail Velocity: instantaneous speed of the tail, derived from the movement of the mean point between the last two tail segments. Angular Velocity: rate of rotational change in the direction of the head, calculated from the angle between vectors connecting the head to a reference body point. Sum of Tail Angles: cumulative angular deviation between consecutive segments along the tail. Mid-Tail Velocity: instantaneous speed of the sixth tail segment, calculated from its displacement over time. Head-Tail Distance: linear distance between the head midpoint and the mean point of the tail’s last segments. Head Acceleration: rate of change of the head’s velocity. Tail Acceleration: rate of change of the tail’s velocity. Angular Acceleration: rate of change in angular velocity. Head Jerk: rate of change of head acceleration. End-Tail Jerk: rate of change of tail acceleration. Sum of Curvature: cumulative discrete triangular curvature of the tail segments. Angle Velocity Heading: angle between the head’s velocity vector and a reference vector from the head toward the body, indicating directional alignment of motion. Curvature Rate: rate of change of tail curvature over time. Note that from each of these parameters the Mean, Maximum, Minimum and/or Range was computed as individual variables used in the dimensionality reduction analysis.

| Variable | Mean | Max | Min | Range | Feature Count |
| --- | --- | --- | --- | --- | --- |
| Head Velocity | X | X |  |  | 2 |
| End-Tail Velocity | X | X |  |  | 2 |
| Angular Velocity | X | X |  |  | 2 |
| Sum of Tail Angles | X | X |  | X | 3 |
| Mid-Tail Velocity | X | X |  |  | 2 |
| Head-Tail Distance | X |  | X |  | 2 |
| Head Acceleration | X | X |  | X | 3 |
| End-Tail Acceleration | X | X |  | X | 3 |
| Angular Acceleration | X | X |  | X | 3 |
| Head Jerk | X | X |  | X | 3 |
| End-Tail Jerk | X | X |  | X | 3 |
| Sum of Curvature | X | X |  | X | 3 |
| Angle Velocity Heading | X | X |  |  | 2 |
| Curvature Rate | X | X |  | X | 3 |
| Total |  |  |  |  | 36 |

